# A Conserved Venous Remodeling Program Governs Inferior Vena Cava Formation in Zebrafish

**DOI:** 10.1101/2025.04.28.650985

**Authors:** Giuseppina Lambiase, Noga Moshe, Calanit Raanan, Rudra N. Das, Karina Yaniv

## Abstract

The Inferior Vena Cava (IVC), the largest venous conduit in mammals, forms through complex remodeling of the embryonic cardinal veins (CVs), a process prone to congenital anomalies, with significant clinical implications. However, the mechanisms underlying IVC formation remain unclear. Here, we identify a conserved IVC in zebrafish that emerges during metamorphosis through remodeling of the embryonic CVs, mirroring the mammalian process. Using in vivo imaging and clonal lineage tracing, we identify the cellular origins and molecular mechanisms controlling IVC formation and demonstrate that the transition from CV to IVC represents a shift from a multifunctional embryonic vein to a specialized adult conduit adapted for high-volume blood return. Overall, our findings illuminate conserved mechanisms of venous remodeling and establish a foundation for investigating congenital venous anomalies.

## Introduction

The cardiovascular system emerges as one of the earliest organs in the embryo, supporting its rapid growth and survival by supplying oxygen and nutrients and removing waste products (*1, 2*). In mammals, including humans, two major venous conduits return deoxygenated blood to the right atrium of the heart. The Superior Vena Cava (SVC) drains the upper body, while the Inferior Vena Cava (IVC) collects blood from the lower extremities and the abdominopelvic region. Interestingly, the IVC does not arise during early embryonic development. Instead, the embryonic cardinal veins (CVs) function transiently before undergoing extensive remodeling and replacement by the IVC.

Around the 4th week of human gestation, a primitive vascular plexus composed of angioblast clusters gives rise to three major sets of veins: the vitelline veins draining the yolk sac, the umbilical veins draining the placenta, and the cardinal veins (CV) draining blood from the embryo’s body (*3*). Between gestational weeks 6th and 8th, the CVs undergo complex remodeling to establish the foundation of the definitive adult venous system, including the **SVC, IVC,** and other major veins. This complex process involves selective regression, anastomosis, and replacement of specific segments of the CVs and their derivatives (*3, 4*). Disruptions in these processes can result in congenital venous anomalies, such as IVC duplication (due to the persistence of right and left supra-cardinal veins), transposition (left-sided IVC), and IVC agenesis (*5–7*), which, although rare and often asymptomatic, can complicate surgical procedures and are associated with increased risk of venous thrombosis and venous stasis (*8*). Therefore, understanding the mechanisms underlying IVC formation is crucial for deciphering the etiology of these malformations and improving their diagnosis and treatment. While the biphasic development of IVC has been documented more than a century ago (*9, 10*), the underlying cellular and molecular mechanisms remain poorly understood. Importantly, it is unclear why this process occurs and what physiological demands drive the replacement of the embryonic CV.

Different animal models have been used to investigate IVC development, including dogs, pigs, sheep, bats, and avians (*11, 12*). Nevertheless, the limited experimental accessibility and precision of cell tracing in these systems have led to at least five different theories describing the origins and mechanisms of IVC formation (*5, 12*). All the proposed models share common elements, such as the early bilaterally symmetric configuration of the embryonic CVs and the anastomosis of the sub-cardinal vein and vitelline veins, which gives rise to the cranial-most portion of IVC. However, the main discrepancy among these models lies in the origin of the caudal part of the IVC (*12*). Over the past decades, the zebrafish has emerged as a valuable model for studying vascular development (*13*). Like humans, zebrafish possess a CV during embryonic stages, which serves as the primary venous vessel carrying deoxygenated blood from the body to the heart (*14, 15*). However, whether zebrafish develop an IVC-like structure during later stages has remained unknown. In this study, we identify the zebrafish IVC and show that it forms during metamorphosis (*16*) through remodeling of the embryonic CV. Using high-resolution longitudinal imaging and endothelial cell-specific clonal lineage tracing, we demonstrate that the IVC originates exclusively from venous endothelial cells and show that the process is regulated by vascular endothelial growth factor A (Vegfa) and its decoy receptor Vegfr1/flt1. Functional analyses reveal that while the embryonic CV serves multiple roles, including macromolecular scavenging (*17, 18*), hematopoietic support (*19*), and harboring specialized angioblasts in its ventral wall (*20–22*), the adult IVC is a streamlined, high-volume conduit. This functional divergence suggests that replacing the CV with the IVC is not merely a structural change but represents a significant vessel specialization event during the organism’s transitions from embryonic to adult stages. Overall, our findings uncover the existence of a well-formed IVC in the adult zebrafish, which develops through the replacement of the CV, mirroring the process observed in mammals, and highlights the zebrafish as a valuable experimentally accessible model for studying the development, function, and related disorders of the IVC. Moreover, this work has implications for understanding similar processes in human IVC formation and for addressing associated congenital anomalies.

## Results

### The axial venous system undergoes extensive remodeling during zebrafish metamorphosis

The anatomical arrangement of the great vessels in the trunk of the early zebrafish larva (i.e., 6 dpf), is well established (*23*). The three large vessels-Dorsal Aorta (DA), Cardinal Vein (CV), and Thoracic Duct (TD), run horizontally and in parallel, spanning the whole trunk (Fig. 1A,B and Fig. S1A,B). Surprisingly, while imaging the posterior trunk of 26 dpf/∼9mm *Tg*(*kdrl:EGFP;lyve1b:dsRed)*(*24*) fish, we noticed the presence of an additional large vessel running between the DA and the CV (Fig. 1C,D and Fig. S1C, magenta dashed line, magenta arrowhead) and wrapped by the TD (Fig. 1D, yellow dashed line and Fig. S1C, white arrowheads). Given its anatomical location resembling the typical position of the IVC in other vertebrates, we hypothesized that this vessel represents the zebrafish equivalent of the mammalian IVC. Hematoxylin and Eosin (H&E) staining of sagittal sections from whole fish at 23 dpf/∼8mm further confirmed the presence of 3 large blood-filled vessels spanning the animal’s trunk (Fig. 1E, DA white dashed lines, CV light-blue dashed lines, IVC magenta dashed lines). Confocal images revealed that the new vessel is also labeled by the venous/lymphatic-specific reporters *lyve1b:dsRed* (Fig. 1D and Fig. S1C, magenta dashed lines) and *mrc1a:EGFP* (Fig. S1D, magenta dashed lines)(*24, 25*), as opposed to the DA that displayed only *kdrl:EGFP* fluorescence (Fig. S1C, white dashed lines), and the TD that was marked only by the *lyve1b* (Fig. 1D, yellow dashed line and Fig. S1C, white arrowhead) and *mrc1a* derived signals (Fig. S1D, yellow dashed line, white arrowhead). Concomitant to the appearance of the new axial vessel, we observed a drastic reduction in the caliber of the CV-while at early stages (13 dpf/∼5mm), the newly formed vessel is narrower than the CV, it undergoes progressive enlargement, with concurrent reduction of the CV (Fig. 1F).

**Fig. 1.**
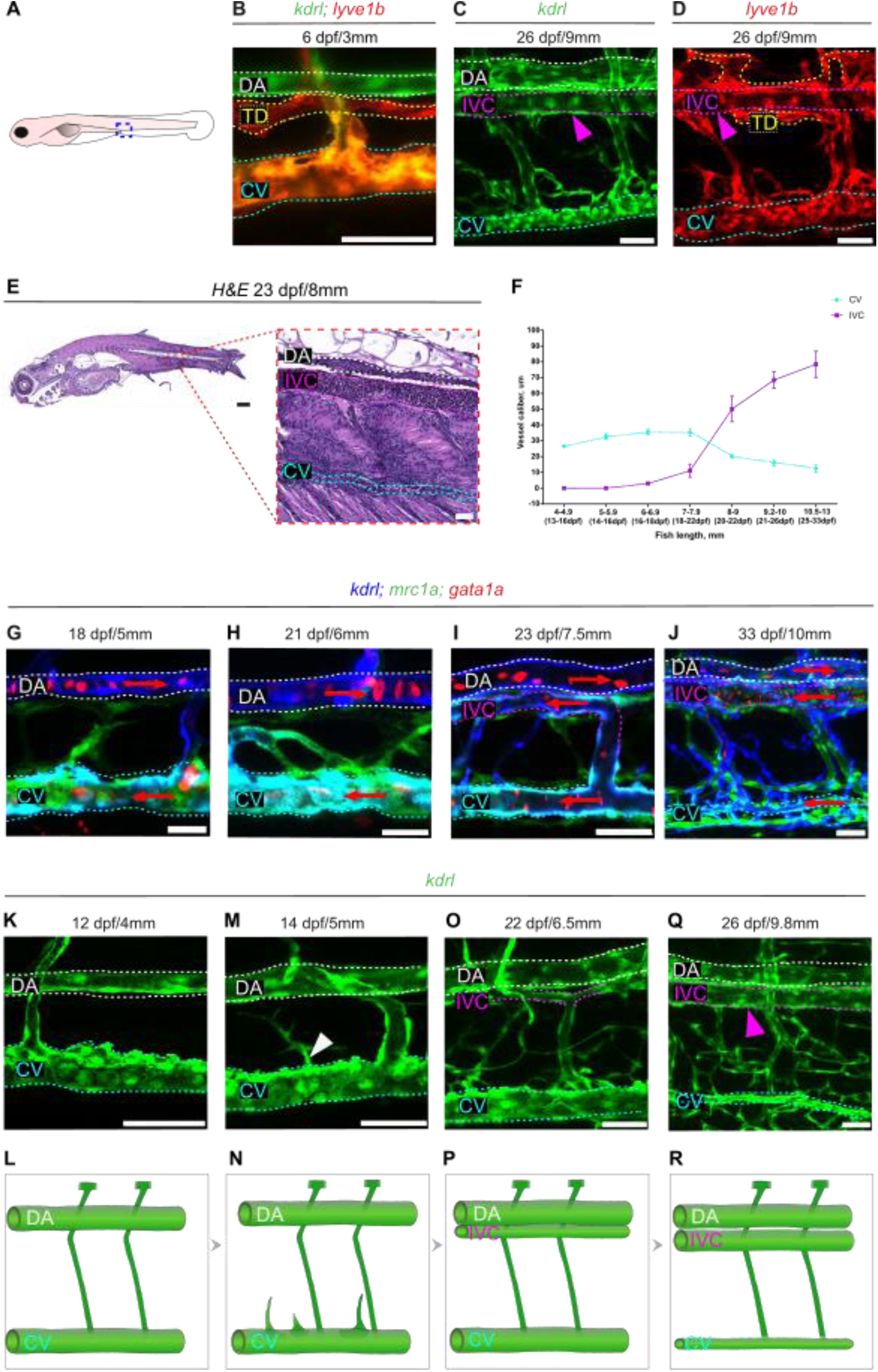
The IVC forms through sequential remodeling of the CV. (**A**) Illustration of zebrafish embryo depicting the area shown in B-D (blue square). (B) Confocal images of Tg(kdrl:GFP;lyve1b:dsRed) trunk vasculature of 6 dpf/3mm larvae, highlighting kdrl+ DA (white dashed lines), kdrl+;lyve1b+ CV (light-blue dashed lines), and lyve1b+ TD (yellow dashed lines). (C) Tg(kdrl:GFP) 26 dpf/9mm fish show the presence of an additional axial blood vessel in the trunk, IVC (magenta dashed lines and arrowhead). (D) At 26 dpf/9mm Tg(lyve1b:dsRed) labels both lymphatics (yellow dashed lines) and the IVC (magenta dashed lines). (E) H&E staining on paraffin sagittal section of 23 dpf/8mm fish, with high magnification of the trunk area highlighting the presence of DA (white dashed lines), CV (light-blue dashed lines), and IVC (magenta dashed lines). (F) Quantification of CV and IVC caliber from 13 dpf/4mm to 33 dpf/13mm fish. (G-J) Erythrocyte flow characterization during IVC development using Tg(kdrl:BFP;mrc1a:GFP;gata1:dsRed) fish. At 18 dpf/5mm and 21 dpf/6mm, erythrocytes are detected only in DA and CV (G,H). At 23 dpf/7.5mm and 33 dpf/10mm IVC also carries erythrocyte flow (I,J). The red arrows depict the direction of flow determined by live imaging. (K,M,O,Q) Progressive formation of the IVC as captured in Tg(kdrl:GFP) fish at 12 dpf/4mm, 14 dpf/5mm, 22 dpf/6.5mm, and 26 dpf/9.8mm. (L,N,P,R) Schematic illustrating the stages of IVC formation. Scale bars, 25 µm (B, C, G, H); 50 µm (C, D, E, I, J, K, M, O, Q). Cardinal vein (CV), light-blue dashed line; Dorsal Aorta (DA), white dashed lines; Thoracic dact (TD), yellow dashed lines; Inferior Vena Cava (IVC), magenta dashed lines.

To further ascertain the identity of the new vessel, we assessed blood flow directionality through live imaging of *Tg*(*gata1a:dsRed)*(*23*) using confocal and light sheet microscopy. We found that at 18 dpf/∼5mm and 21 dpf/∼6mm the DA carries blood from the heart to the trunk, while the CV brings blood back to the heart (Fig. 1G,H and movie S1). Later, at 23 dpf/∼7.5mm, as the new vessel becomes lumenized and patent, erythrocyte flow is observed toward the heart, suggesting it is of venous nature (Fig. 1I and movie S2). At this stage, the CV and the newly formed IVC share the venous blood flow. However, as the IVC continues to expand, a significant increase in the proportion of erythrocytes routed through it is observed, along with a corresponding reduction in flow through the much-thinned CV (Fig. 1J and movie S2).

We then set out to characterize the steps leading to the establishment of the IVC. We began by imaging the caudal trunk of *Tg*(*kdrl:EGFP;lyve1b:dsRed)* animals at 12 dpf/∼4mm, where we clearly detected the DA, CV, and TD (Fig. 1K,L and Fig. S1E,F). Later on, at 14 dpf/∼5mm, we observed CV-derived sprouts extending dorsally (Fig. 1M,N and Fig. S1G,H white arrowhead). By 22 dpf/∼6.5mm the sprouts reach the level of the DA where they bifurcate and anastomose to generate a new lumenized vessel (Fig. 1O,P and Fig. S1I,J, magenta dashed lines) that expands by ∼26 dpf/∼ 9.8mm (Fig. 1Q,R and Fig. S1K,L, magenta dashed line and magenta arrowhead). As the IVC expands, we observed a concurrent reduction of the CV caliber over time (Fig. 1F, K-R, light-blue dashed lines). Finally, we examined histological sections of adult zebrafish to determine if the emerging IVC persists through adulthood. Indeed, we detected a major vein running ventral and parallel to the DA (Fig. S1M, magenta dashed lines, magenta arrowhead), whereas no remnants of the CV were observed at this stage (Fig. S1M, light blue asterisk). Together, our results demonstrate that the embryonic-born CV undergoes extensive remodeling and regression while a juvenile-born vein sets in ventral to the DA. The striking similarity between this process and the one described during mammalian IVC development led us to conclude that the newly formed vessel represents the zebrafish equivalent to the mammalian IVC, which takes over the main function of the CV, i.e., bringing deoxygenated blood back to the heart. Moreover, our data show that the IVC forms during metamorphosis (i.e., post-embryonic development) and persists till adulthood, replacing the preexisting CV.

### Cellular Origins of the IVC

To investigate the ontogeny of the IVC, we utilized our previously established EC-specific *brainbow*-based clonal analysis tool – *flibow* (*24*), in combination with Tg(*kdrl:CreER^T2^*)(*26*), to specifically label blood ECs. We first asked whether the major axial vessels contribute to the formation of the IVC. Treating 1.5, 2, 2.5, and 3 dpf *Tg(kdrl:CreER^T2^;flibow)* embryos with 4-hydroxytamoxifen (4-OHT) resulted in efficient mosaic labeling of the CV and its derivatives. In contrast, the DA was rarely labeled (Fig. S2A-D and Table 1), enabling us to distinguish potential contributions from these two vessels. Following 4-OHT administration, embryos were raised under normal conditions and screened for the presence of sparse and distinguishable labeled EC clones in the CV and DA between 5-10 dpf, i.e., before the initiation of IVC formation. The selected animals were then imaged at multiple time points for robust and reliable tracing of the identified clones (Fig. S2E). We found that out of all *larvae* carrying labeled CV-ECs (n=28), 93% showed clonally related progeny in the IVC (Table 1). For instance, in fish #1 (Fig. 2A), we identified two distinctly labeled ECs at 14 dpf/∼5mm-one located on the dorsal (Fig. 2A, yellow EC, magenta arrowhead) and one on the ventral (Fig. 2A, cyan EC, white arrowhead) side of the CV. Imaging of the same larva at 21 dpf/∼6.2mm and 29dpf/∼9mm (Fig. 2B,C, magenta arrowhead) revealed that only the dorsal EC generated clonally related progeny, which contributed to the IVC. In contrast, ECs located in the ventral CV did not contribute to IVC formation and either remained stable throughout the duration of the experiment or sprouted ventrally to generate other vessels (Fig. 2A-C, white arrowhead). In a cohort of 30 fish analyzed for dorsal and ventral clone location, ∼63% of the labeled dorsal ECs contributed to the IVC, whereas only ∼29% of ventral ECs did (Fig. S2F). These data are in line with previous reports demonstrating the angiogenic properties of dorsal CV-ECs (*27–29*), while highlighting the more progenitor nature of ECs located on the ventral wall of the CV (*20, 22*). Similar results were obtained following imaging of fish #2 at 14 dpf/∼5mm, 16 dpf/∼6mm, and 20 dpf/∼8.5mm (Fig. 2D-F, magenta arrowheads) and fish #3 at 12 dpf/∼4.5mm, 16 dpf/∼5.5mm, 22 dpf/∼7mm and 24 dpf/∼9mm (Fig. S2G-J, magenta arrowheads).

**Fig. 2.**
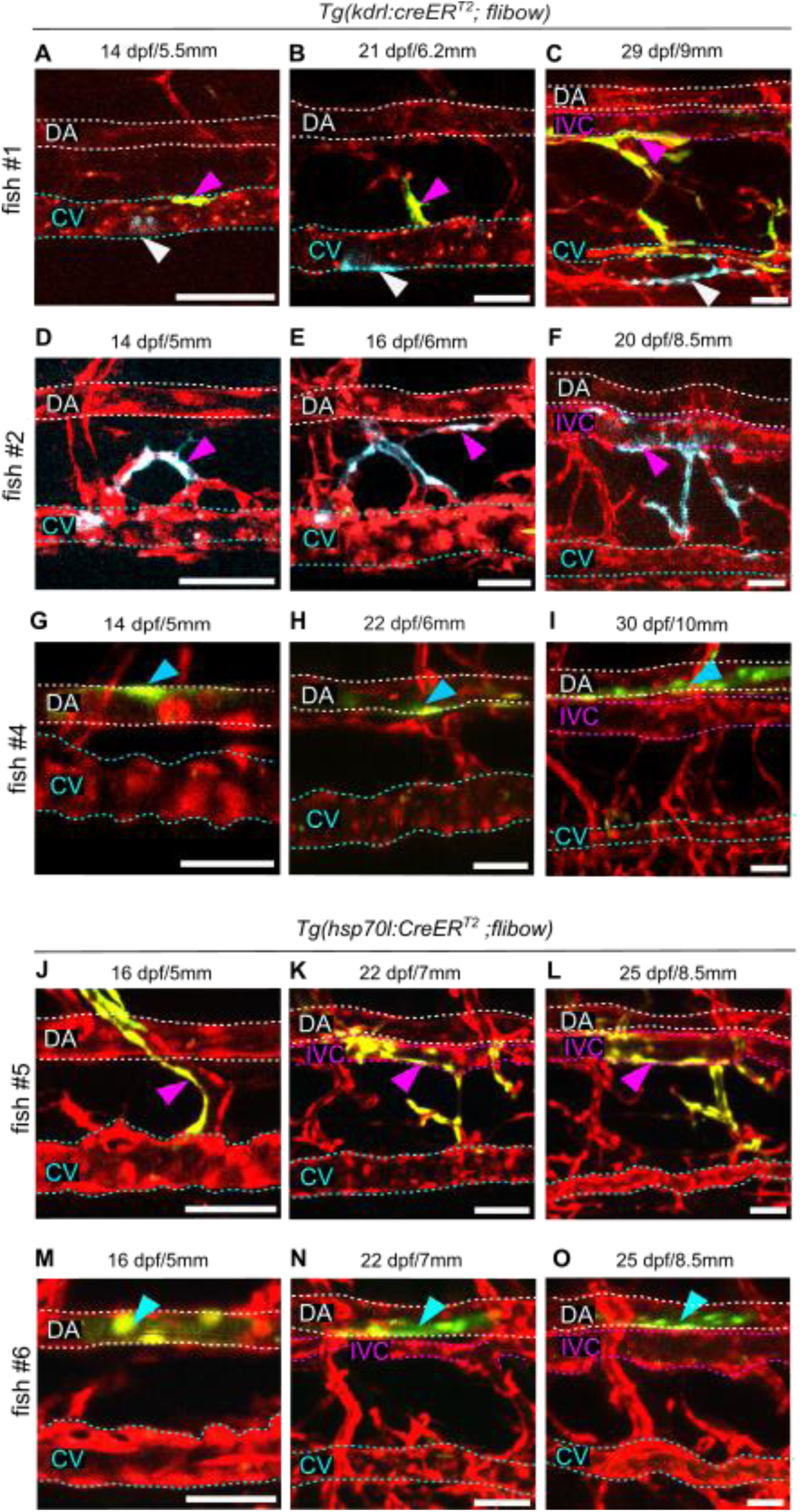
Clonal analysis reveals the venous origins of the IVC. (**A-I**) Reiterative imaging of three individual Tg(kdrl:CreER^T2^;flibow) animals over time. (A-C) Confocal images of Fish#1 at 14 dpf/5.5mm, 21 dpf/6.2mm and 29 dpf/9mm. An EC at the dorsal CV (yellow cell, magenta arrowhead) gives rise contributes to the IVC (**C**), while a ventral CV-EC (cyan cell, white arrowhead) gives rise to other vessels. (D-F) Fish#2 imaged at 14dpf/5mm, 16dpf/6mm and 20dpf/8.5mm shows a clone progressing from the CV sprout (D, magenta arrowhead) to the nascent (E) and developed (F) IVC. (G-I) Fish#4 imaged at 14 dpf/5mm, 22 dpf/6mm and 20 dpf/8.5mm shows a DA-derived clone (green cell, cyan arrow) that does not contribute to the IVC. (J-O) Reiterative imaging of two different Tg(hsp70l:CreER^T2^;flibow) animals over time (J-L) Fish#5 imaged at 16 dpf/5mm, 22 dpf/7mm and 25 dpf/8.5mm shows CV-derived clones in (J, yellow cells, magenta arrowhead) contributing to the IVC (K,L). (M-O) Fish#6 shows DA clones (M, yellow cells, cyan arrowhead) that are not detected in the IVC at later stages (N,O). Scale bars 50 µm (A-N). Cardinal vein (CV), light-blue dashed lines; Dorsal Aorta (DA), white dashed lines; Inferior Vena Cava (IVC), magenta dashed lines.

**Table 1.** List of kdrl:creER^T2^;flibow and hsp70l:creER^T2^;flibow animals traced for lineage analyses. Information on the induction conditions, stages that were imaged, and EC subsets labeled.

Unlike the CV, we never found clones derived from the DA populating the IVC (fish #4; Fig. 2G-I, light-blue arrowhead), indicating that this major vein is established primarily via sprouting of preexisting CV-ECs and their derivatives. These observations were further confirmed through an unbiased lineage tracing strategy involving *Tg(hsp70l:CreER^T2^)* animals, where CreER^T2^ is ubiquitously expressed, while the *brainbow* construct is specifically expressed in ECs upon heat and 4-OHT activation (*24*). In animals treated at 3dpf, only venous-derived ECs were found in the IVC (fish #5; Fig. 2J-L, magenta arrowheads) with no contribution from DA-ECs (fish #6; Fig. 2M-O, light-blue arrowhead).

### Molecular control of IVC formation

Since venous sprouting in the embryo is known to be controlled by VegfC/VegfD signaling via the VegfR3/Flt4 receptor (*27, 28, 30*), we first interrogated the involvement of this pathway in IVC formation. Given that homozygous *vegfc* mutants are embryonic lethal (*31*), we examined *vegfd* and *flt4* mutants, which live through adulthood (*24, 32*). We found that *in Tg(kdrl:GFP;vegfd^-/-^),* both venous sprouting (Fig. S3A-B) and IVC formation (Fig. S3C,D, magenta dashed lines) proceeded normally, with no significant differences in IVC caliber (Fig. S3E). Similar results were obtained following analysis of *flt4^-/-^* animals, which also possess a normal IVC (Fig. S3F-H, magenta dashed lines). Since *flt4* mutants completely lack trunk lymphatics (Fig. S3I,J, yellow asterisks) (*24, 27*), our data indicate that the neighboring lymphatic vessels are not required for proper IVC formation and do not contribute ECs to this structure, further confirming our lineage tracing results.

We then assessed the involvement of VegfA signaling in IVC formation by analyzing *vegfaa* and *vegfab* mutants. Previous studies established that *vegfaa^-/-^* embryos feature complete loss of the DA and expansion of the CV, and die around 3 dpf (*33, 34*). By contrast, *vegfab* mutants show only mild vascular defects and survive through adulthood (*34*). Accordingly, double homozygous mutants *(vegfaa^-/-^;vegfab^-/-^)* are embryonic lethal (*33*), whereas all other allelic combinations are viable through adult stages. We focused our analyses primarily on *vegfaa^+/-^;vegfab^-/-^* fish, which possess only one wt allele. During the initial sprouting phase, between 5-6mm, *vegfaa^+/-^;vegfab^-/-^*fish showed a visible reduction in the number of sprouts emerging from the CV as compared to wt siblings (Fig. 3A,B, white arrowhead, C). This resulted in the formation of a truncated and abnormal IVC, with several segments missing, at later stages (*i.e.,* at ∼8-10mm) (Fig. 3D,E). To better characterize the phenotype, we used a major shunt connecting the CV and the IVC in both wt and mutant animals as a landmark to divide the trunk into rostral and caudal regions (Fig. 3E-H). We found that *vegfaa^+/-^;vegfab^-/-^* fish exhibited a lumenized albeit slightly narrower IVC in the rostral trunk (Fig. 3E,H, magenta dashed lines, magenta arrowheads, I), accompanied by a normally regressing CV (Fig. 3E,H, light blue arrowheads, J). In contrast, the caudal trunk showed a thin and fragmented IVC (Fig. 3E,H, yellow arrowheads, magenta asterisks, K,L) and a larger CV (Fig. 3E,H, blue arrowhead, light-blue dashed lines, M). These results hint at a putative compensatory mechanism wherein CV regression is intrinsically linked to IVC formation, with the CV potentially maintaining its structure when IVC development is compromised.

**Fig. 3.**
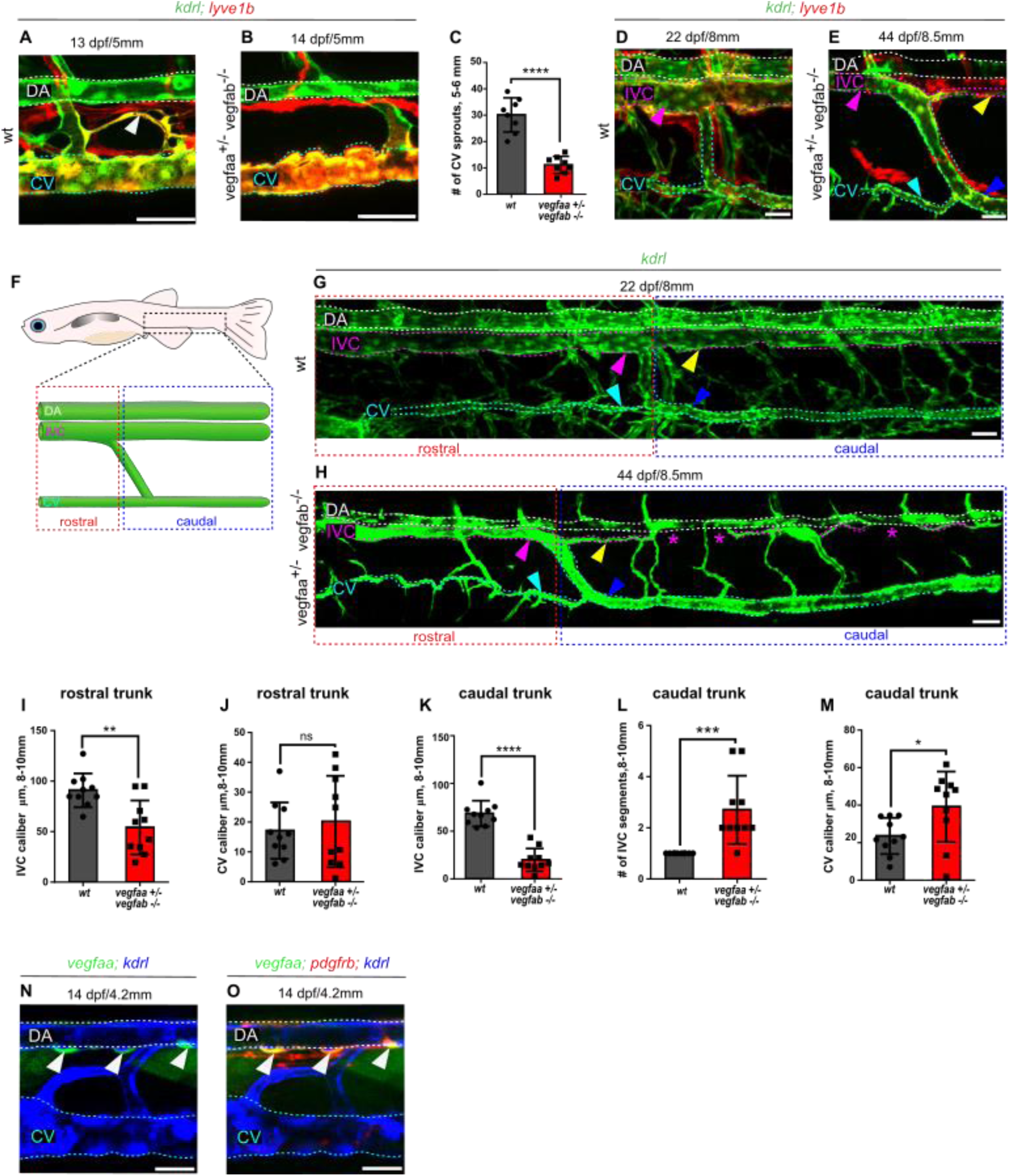
Vegfa signaling is required for CV sprouting and proper IVC morphogenesis. (**A-B**) Confocal images of Tg(kdrl:EGFP lyve1b:dsRed) wt larvae at 13 dpf/5mm show CV-emerging sprouts (A, white arrowhead) that are nearly absent in 14 dpf/5mm vegfaa^+/-^;vegfab^-/-^ siblings (B). (C) Significantly reduced number of sprouts in vegfaa^+/-^;vegfab^-/-^ (5-6mm size) larvae compared to wt siblings (n=8, t-test, p < 0.05). (D) Tg(kdrl:EGFP;lyve1b:dsRed) wt fish at 22 dpf/8mm show a well-formed IVC (magenta arrowhead) and a narrow CV (light-blue dashed lines). (E) Confocal image of Tg(kdrl:EGFP;lyve1b:dsRed); vegfaa⁺/⁻ vegfab⁻/⁻ fish at 44 dpf/8.5 mm showing segment-specific phenotypic differences within the mutant trunk. In the rostral trunk, the IVC is properly lumenized (magenta arrowhead) and the CV is narrowed (cyan arrowhead), while in the caudal trunk, the IVC appears thin (yellow arrowhead) and the CV remains abnormally wide. (F) Schematic of the larval trunk illustrating an anatomical shunt (white asterisk) that connects the CV and IVC used to divide the trunk into two segments for analysis: the rostral trunk (red dashed box), and the caudal trunk (blue dashed box). (G) Confocal image of a 22 dpf/8mm Tg(kdrl:GFP) wt fish showing CV regression (light-blue and blue arrowheads) and intact IVC (magenta and yellow arrowhead) in both areas. (H) Trunk vasculature of 44 dpf/8.5mm Tg(kdrl:GFP; vegfaa^+/-^;vegfab ^-/-^) animal showing normal rostral IVC (magenta arrowhead) and a normal regression of CV (light blue arrowhead) but a thin and fragmentated caudal IVC (yellow arrowhead, magenta asterisks) and persistent CV (blue arrowhead). (I) Rostral IVC caliber at 8-10mm is reduced in vegfaa^+/-^;vegfab^-/-^ compared to wt (n=10,t-test, p < 0.005). (J-M) Quantification of CV caliber size in rostral and caudal regions. (J) Rostral CV caliber is unchanged in vegfaa^+/-^;vegfab^-/-^ mutants (n=10, t-test, p>0.05). (K) Caudal IVC caliber is significantly reduced in vegfaa^+/-^;vegfab^-/-^ mutants (n=10, t-test, p< 0.05). (L) Th number of IVC fragments is significantly increased in the caudal trunk of vegfaa^+/-^;vegfab^-/-^ (n=10, t-test, p<0.05). (M) CV caliber is wider in the caudal trunk of vegfaa^+/-^;vegfab^-/-^ animals (n=10, t-test, p >0.05). (N, O) vegfaa:EGFP expression at 14 dpf/5.4 mm localizes to perivascular cells (white arrowhead), overlapping with Tg(pdgfrb:Gal4; uas:NTR-mCherry) expression (O). Scale bars 50 µm (A,B,D,E,N,O), 100 µm (G,H). Cardinal vein (CV), light-blue dashed lines; Dorsal Aorta (DA), white dashed lines; Inferior Vena Cava (IVC), magenta dashed lines.

Analysis of all other *vegfaa* and *vegfab* mutant combinations, i.e., *vegfaa^+/-^; vegfab ^+/-^* (Fig. S3K,L)*, vegfaa^+/-^;vegfab*^+/+^ (Fig. S3M) and *vegfaa^+/+^; vegfab* ^-/-^ (Fig. S3N) at 14 dpf/5mm did not show any alterations in the number of CV sprouts (Fig. S3O-Q) or the resulting IVC (Fig. S3R-U). Finally, we interrogated the source of Vegfaa during IVC formation. At 14 dpf/4.2mm, right before the onset of CV sprouting, Vegfaa was present in *pdgfrb+* pericytes surrounding the DA (Fig. 3N,O, white arrowheads), suggesting it might be released by pericytes to attract CV sprouts forming the IVC formation.

Altogether, these data show that pericyte-derived Vegfa signaling is specifically required for IVC formation, and although the allelic combination (*vegfaa^+/-^;vegfab^-/-^*) does allow for blood vessels to develop in the early fish embryo, structures like the IVC, which form during metamorphosis are particularly sensitive to VegfA alterations.

Various reports have highlighted the involvement of Vegfr1/flt1 in certain settings of venous sprouting (*35–38*). For instance, during the formation of the caudal fin vasculature, venous sprouts generating arteries activated the arterial marker *flt1* in a VEGFA-dependent manner (*36*). Similarly, *flt1* was found to be expressed in brain venous sprouts emerging from the PHBC, that give rise to the Central Arteries (CtAs) (*35*). We therefore wondered whether a similar mechanism could account for IVC formation in response to VegfA signaling. Indeed, at 15 dpf/∼5mm we detected clear expression of the *Tg(flt1^BAC^:YFP)* (*39*) reporter not only in the DA (Fig. 4A,B), but also in dorsal ECs sprouting from the CV (Fig. 4A,B, white arrowhead), as well as in the nascent IVC (Fig. 4C,D, magenta dashed line) at 22dpf/∼8mm. Similarly to *flt1,* also *dll4* is reported to be expressed in the tip cells during the artery formation (*40, 41*). Indeed, we detected expression of the Notch ligand *dll4* in the initial CV sprouts (Fig. 4E,F, white arrowhead) as well as in the IVC at 18 dpf/∼8mm (Fig. 4G,H, magenta dashed lines). Taken together, our results indicate that, similar to previously described cases of venous arterialization (*36, 39*), the sprouts that give rise to the IVC are regulated by the *vegfa/flt1/dll4* axis. Although these sprouts retain venous identity and ultimately form a vein, they transiently activate signaling pathways typically associated with arterial tip-cell behavior.

**Fig. 4.**
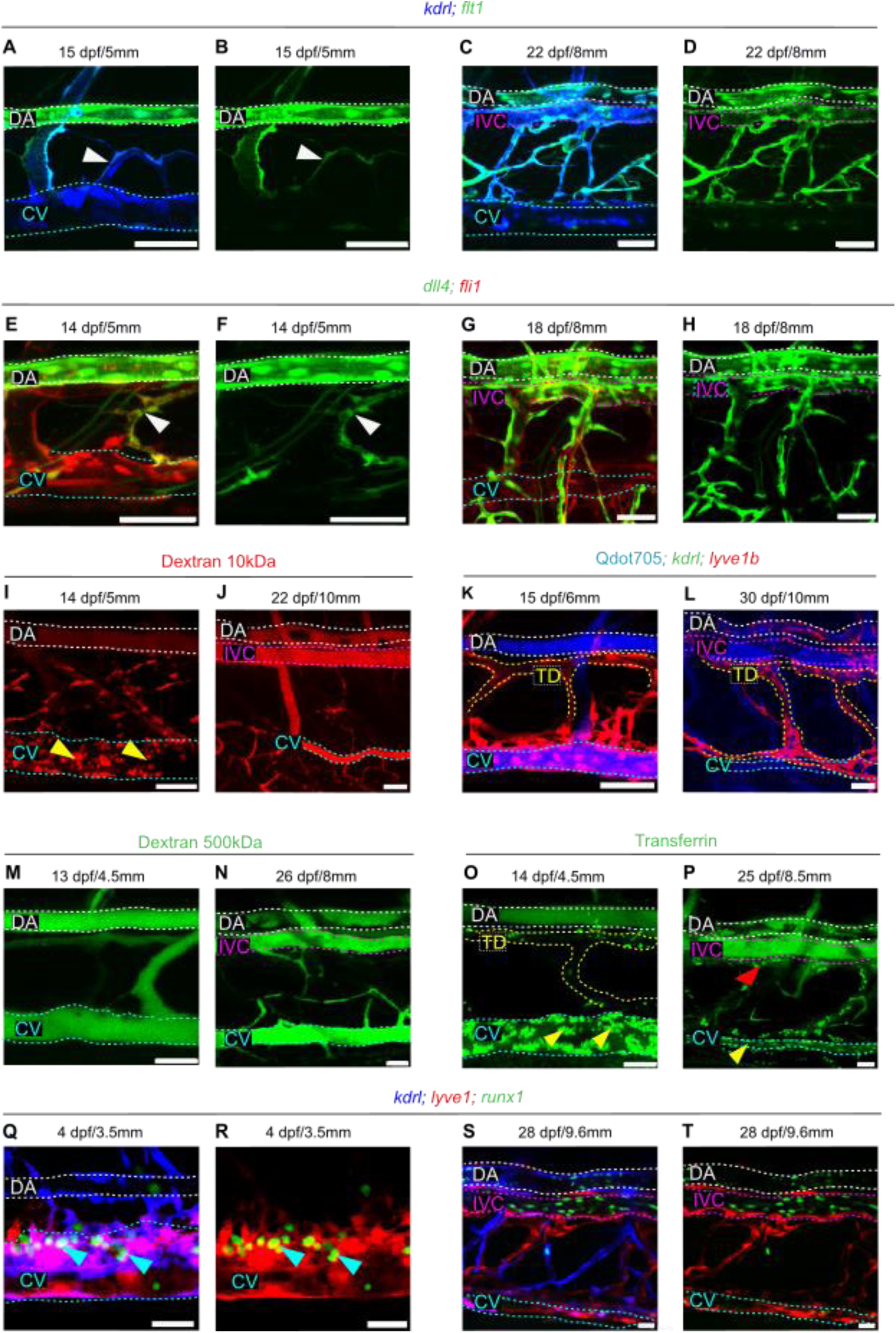
Functional divergence between the embryonic CV and the adult IVC. (**A-D**) Tg^BAC^(flt1:YFP);Tg(kdrl:BFP) trunk of 15 dpf/5mm and 22 dpf/8mm fish show flt1 expression in CV sprouts (white arrowhead) and IVC (magenta dashed lines), indicating shared molecular features. (E-H) Tg(dll4:GAL4,UAS:GFP;fli1:dsRed) trunk of 14 dpf/5mm and 18 dpf/8mm fish showing similar dll4 expression in CV sprouts (white arrowhead) and IVC (magenta dashed lines). (I–P) Angiography using different tracers reveals a developmental loss of EC scavenger function from the CV to the IVC. (I, J) Rhodamine Dextran 10kDa is readily internalized by CV-ECs at 14 dpf/5mm (yellow arrowheads), but no uptake is detected in either CV (light-blue dashed lines) or IVC (magenta dashed lines) by 22 dpf/10mm. (K,L) Qdot705 particles are not internalized by CV or IVC at either 15 dpf/6mm or 30 dpf/10mm, indicating size-restricted uptake.(M,N) Similarly, no uptake is observed with high-molecular-weight FITC-Dextran (500kDa) at 13 dpf/4.5mm and 26 dpf/8mm. (O) Transferrin-488 is internalized by CV-ECs and TD at 14 dpf/4.5mm (yellow arrowheads). (P) At 25 dpf/8.5mm, this uptake persists only in the CV (yellow arrowhead) and TD (red arrowhead) while the IVC remains negative (magenta dashed lines). (Q**-**R) Tg(kdrl:BFP;lye1:dsRed;runx1:GPF) fish at 4 dpf/3.5mm shows runx1+ cells within the CV-EC pockets (light-blue arrowheads) and in circulation. (S,T) At 28 dpf/9.6 mm, runx1+ cells are detected in circulation but not wrapped by IVC-ECs in Tg(kdrl:BFP;lyve1:dsRed;runx1:GPF). Scale bars 50 µm (A-T). Cardinal vein (CV), light-blue dashed lines; Dorsal Aorta (DA), white dashed lines; Inferior Vena Cava (IVC), magenta dashed lines.

### CV and IVC have distinct functional properties

Having established the developmental trajectory of the IVC, we next investigated the physiological significance of replacing the CV with the IVC. Specifically, we asked whether these two vessels serve different functions. In addition to transporting deoxygenated blood to the heart, the embryonic CV has been shown to contain ECs displaying regional specialization and varying functionality. For instance, the floor of the CV homes a population of specialized angioblasts, which give rise to several EC types, including arteries, veins, and lymphatics (*20, 22*). Likewise, ECs in the dorsal side of the CV are more “angiogenic” and sprout in response to VegfA/VegfC signals (*28, 29, 42, 43*). Toward the tail, the caudal segment of the CV, known as caudal hematopoietic tissue (CHT), serves as a niche for hematopoietic stem cell expansion (*19, 44*). Finally, a specialized “scavenger” EC type, which displays high endocytic capabilities, has also been identified in the embryonic CV (*18*). Given that the CV represents the main source of IVC-ECs, we decided to investigate whether some of the functions of CV-ECs are maintained in the IVC, with a particular focus on the scavenging function. CV-ECs were shown to uptake a variety of substrates, such as lipoproteins, glycosaminoglycan proteins, and polysaccharides at 5 dpf. In addition, anionic nanoparticles between 50 and 250 nm were shown to be taken up by CV-ECs, solely dependent on surface charge (*18*). Indeed, intravascular injection of Rhodamine Dextran 10kDa at 14 dpf/5mm revealed robust uptake by ECs in the CV, particularly in its caudal region (Fig. 4I, yellow arrowheads). In contrast, neither the newly formed IVC nor the residual CV exhibited scavenging activity at 22 dpf/∼10mm (Fig. 4J), indicating that despite a shared origin, IVC ECs lack the scavenger function characteristic of their CV counterparts, reflecting a shift in EC specialization during venous remodeling. Large-sized particles, such as Qdot705 (Fig. 4K,L) or Dextran 500kDa (Fig. 4M,N), were not internalized by either the CV or the IVC at both early and late stages.

Next, we evaluated the contribution of receptor-mediated endocytosis to the scavenging properties of CV-ECs and whether they change as the IVC is formed. To this end, we performed intravascular injection of Transferrin (Trf-488), an iron-binding protein that binds the transferrin receptor and enters the cell through receptor-mediated endocytosis (*45*). As seen in Fig. 4O, Trf-488 was rapidly internalized by ECs of the CV (yellow arrowheads) and by LECs of the TD at 14 dpf/∼4.5mm (Fig. 4O, yellow dashed lines) and 25 dpf/∼8.5mm (Fig. 4P, light blue dashed lines, yellow and red arrowheads), but not by ECs of the IVC at similar stages (Fig. 4P, magenta dashed lines).

Finally, along with their scavenger function, embryonic CV-ECs in the CHT are reported to function as a niche supporting hematopoietic stem cell expansion and differentiation (*19, 44*). Through imaging of *Tg(kdrl:BFP;lyve1b:dsRed;runx1:GFP)* (*46*), we found that while *runx1*+ cells were detected within EC pockets in the CV-CHT at 4 dpf/∼3.5mm (Fig. 4Q,R light blue arrowheads; movie S3), the IVC does not harbor HSCs at 28 dpf/9.6mm (Fig. 4S,T; movie S4). Altogether, our findings reveal that the juvenile CV serves multiple distinct functions, which are not shared by the IVC, underscoring the functional differences between these two veins.

Overall our results uncover a previously unrecognized program of venous remodeling during zebrafish metamorphosis marked by the regression of the CV and its replacement by the IVC. This process is driven exclusively by venous ECs with no contribution from arterial or lymphatic lineages. We further show that IVC formation critically depends on Vegfaa signaling, while Vegfd and Flt4-key regulators of embryonic venous and lymphatic sprouting are not involved in this process. Interestingly, although the sprouts forming the IVC retain venous identity, they transiently express arterial-associated markers such as Flt1 and Dll4, indicating co-option of tip-cell-like programs during vein-to-vein remodeling. Functionally, this transition reflects a shift from the embryonic CV’s diverse roles to the IVC’s specialization in efficient high-volume blood transport. Together, these findings reveal a conserved and mechanistically distinct program of venous remodeling and establish zebrafish as a powerful model for studying IVC development and its associated anomalies.

## Discussion

The Inferior Vena Cava (IVC) is the primary venous conduit for returning deoxygenated blood from the lower body to the heart in mammals. Unlike the other major vessels, the Dorsal Aorta and the Thoracic Duct, which emerge during early embryogenesis, the IVC forms through a biphasic process involving extensive remodeling of embryonic cardinal veins (CVs). This complex ontogeny contributes to congenital IVC anomalies that appear in 0.3% to 10.14% of the population (*47*). When undiagnosed, these defects can complicate surgeries and can be a risk factor for deep venous thrombosis as a result of inadequate venous drainage of the lower extremities through collateral circulation (*8*).

Although the biphasic development of IVC has been recognized for over a century, its cellular and molecular basis has remained elusive, largely due to the inaccessibility of post-embryonic stages in large animal models, which have provided only partial and often contradictory conclusions (*12*). In this study, we demonstrate that zebrafish undergo a conserved biphasic process of CV remodeling, resulting in the formation of an IVC-equivalent vessel, providing an experimentally amenable model system to investigate questions pertinent to IVC development and function.

While the development of the cardinal venous system during embryonic stages is well described in zebrafish (*48*), little is known about the vascular changes that occur during metamorphosis, which help stabilize the adult vasculature. We show that the zebrafish IVC forms during larva-to-juvenile transition and replaces the embryonic CV as the main venous conduit. Anatomically, the IVC runs ventral and parallel to the DA, mirroring its mammalian counterpart. The newly identified zebrafish vessel brings deoxygenated blood from the caudal areas, is located parallel to the DA across the trunk, and features a developmental process that closely mirrors the formation of the mammalian IVC. These anatomical and developmental similarities strongly support the notion that this vessel is the zebrafish equivalent of the mammalian IVC.

Revealing the cellular origins of newly formed vessels is important as it provides important clues on the type of cellular transformation and the underlying signaling mechanisms. In addition, it has been shown that the origins help determine the functional properties of vessels (*24*). Nevertheless, tracing progenitors in vertebrate postembryonic development is a major challenge due to technical difficulties, and requires specialized techniques (*49*). In the absence of robust lineage tracing model systems enabling the study of the IVC, the literature has been confounded by various reports from humans and large mammals.

In this work, we leverage a multicolor EC lineage tracing *(“flibow”)* in conjuction with reiterative live imaging, to trace the origins of the IVC, and investigate the contributions of different vascular beds to IVC development. Our data reveal that the newly formed IVC arises exclusively from venous sources, with no contributions from arterial and lymphatic EC progenitors, indicating that IVC formation doesn’t require major EC identity changes.

Surprisingly, however, we find that this process of “venous-to-venous” sprouting is not controlled by the Vegfc/Vegfd/Flt4 signaling pathway but makes use of arterial/tip cell sprouting mechanisms. Indeed, our findings highlight the pivotal role of the Vegfa signaling pathway (*50*) and pinpoint the importance of the two zebrafish Vegfa paralogs, Vegfaa and Vegfab. Through analysis of various *vegfaa* and *vegfab* mutant combinations, we show that a single functional copy of *vegfaa* is sufficient to support early vascular development, but not adequate to sustain proper IVC formation in the absence of *vegfab*. This indicates that while *vegfaa* plays a dominant role, *vegfab* contributes a supportive function that becomes essential under reduced *vegfaa* dosage. In contrast, single *vegfab* mutants display no overt phenotype, highlighting a degree of functional redundancy during early development. These results underscore a context-dependent requirement for Vegfa paralogs during post-embryonic vascular remodeling and point to the IVC as a particularly sensitive structure that relies on precise regulation of Vegfa dosage.

We also detect expression of *flt1* and *dll4*—genes typically associated with arterial endothelium and tip cell behavior, within ECs sprouting dorsally from the CV. Notably, these sprouts do not acquire arterial identity, as shown by our lineage tracing. Instead, the expression of these markers appears to be transient and localized to actively sprouting venous cells, suggesting that elements of the arterial tip-cell transcriptional program are co-opted during this phase of venous remodeling, consistent with similar findings in other vascular contexts (*51, 52*). In the case of IVC formation, however, the sprouts retain their venous identity and ultimately give rise to a vein. This points to a distinct phenomenon where tip-cell machinery is utilized not for fate switching, but for structural remodeling within the venous compartment. Taken together, our data support a model in which localized, pericyte-derived Vegfa activates a *vegfaa/flt1/dll4* axis that drives precise and directional sprouting from the CV, employing a tip-cell-like transcriptional program to orchestrate vein-to-vein remodeling. This occurs independently of the canonical Vegfc/Flt4 pathway that governs embryonic venous and lymphatic sprouting (*27, 42, 43*), emphasizing that post-embryonic vascular remodeling engages distinct and spatially restricted signaling cues.

We also uncover functional divergence between the CV and IVC. While the embryonic CV supports diverse physiological roles, including macromolecule scavenging (*18*) and hematopoietic stem cell expansion (*46, 53*), the IVC appears structurally simpler and specialized for high-volume blood transport. Our tracer uptake assays confirm that IVC ECs do not retain the scavenging capacity of their CV progenitors. These differences suggest that the transition from CV to IVC is driven not only by anatomical remodeling but also by a shift in EC specialization, enabling the circulation system to meet the increased hemodynamic demands of the growing organism.

The timing of IVC development in fish coincides with metamorphosis a period marked by rapid growth and increased circulatory load (*54*). Unlike the DA and TD, which form during embryogenesis and persist into adulthood, the CV appears to be insufficient to support the needs of the juvenile and adult vasculature. This may relate to the sinusoidal-like nature of the CV endothelium (*18*), which is optimized for exchange but poorly suited for efficient bulk flow. The emergence of the IVC thus reflects a functional adaptation, transforming a multifunctional embryonic vessel into a streamlined adult conduit optimized for high-throughput venous return. In conclusion, our study reveals a conserved, biphasic mode of IVC development in zebrafish and uncovers the cellular and molecular mechanisms governing this process during post-embryonic growth. These data not only advance our understanding of vertebrate vascular remodeling but also provide a genetically tractable system to investigate congenital venous anomalies and the physiological regulation of vein identity.

## Acknowledgments

The authors thank H. Raviv and Y. Yogev (Weizmann Institute, Israel) for technical assistance; G. Almong, R. Hofi, E. Glozman, D. Zuberi, E. Regev and R. Brihon (Weizmann Institute, Israel) for superb animal care, Y. Addadi (Weizmann Institute, Israel) and DST-FIST Confocal Microscope Facility at SNIoE for assistance with imaging experiments. The authors are grateful to all the members of the Yaniv lab for discussion, technical assistance, critical reading of the manuscript, and continuous support. K.Y. is the incumbent of the Enid Barden and Aaron J. Jade Professorial Chair in Memory of Canter John Y. Jade. RND is supported by core funding from SNIoE.

## Funding

ERC CoG (LymphMap 818858) (KY)

Minerva Foundation 712610 (KY)

H&M Kimmel Institute for Stem Cell Research (KY).

Weizmann SABRA – Yeda-Sela – WRC Program (KY)

EMBO long-term fellowship (ALTF 1532-2015) (RND)

Edith and Edward F. Anixter Postdoctoral Fellowship, Weizmann Institute of Science (RND)

Senior postdoctoral fellowship, Weizmann Institute of Science (RND)

Grant No. SR/FST/LS-1/2017/59(c) (RND)

Lombroso Fellowship (GL)

Estate of Olga Klein Astrachan and the Estate of Mady Dukler (NM)

## Author contributions

Conceptualization: RND, KY.

Methodology: GL, NM, RND

Investigation: GL, RND, KY.

Visualization: GL, RND, KY.

Funding acquisition: KY.

Project administration: KY.

Supervision: RND, KY

Writing – original draft: GL, RND, KY.

---

**Fig. S1.**
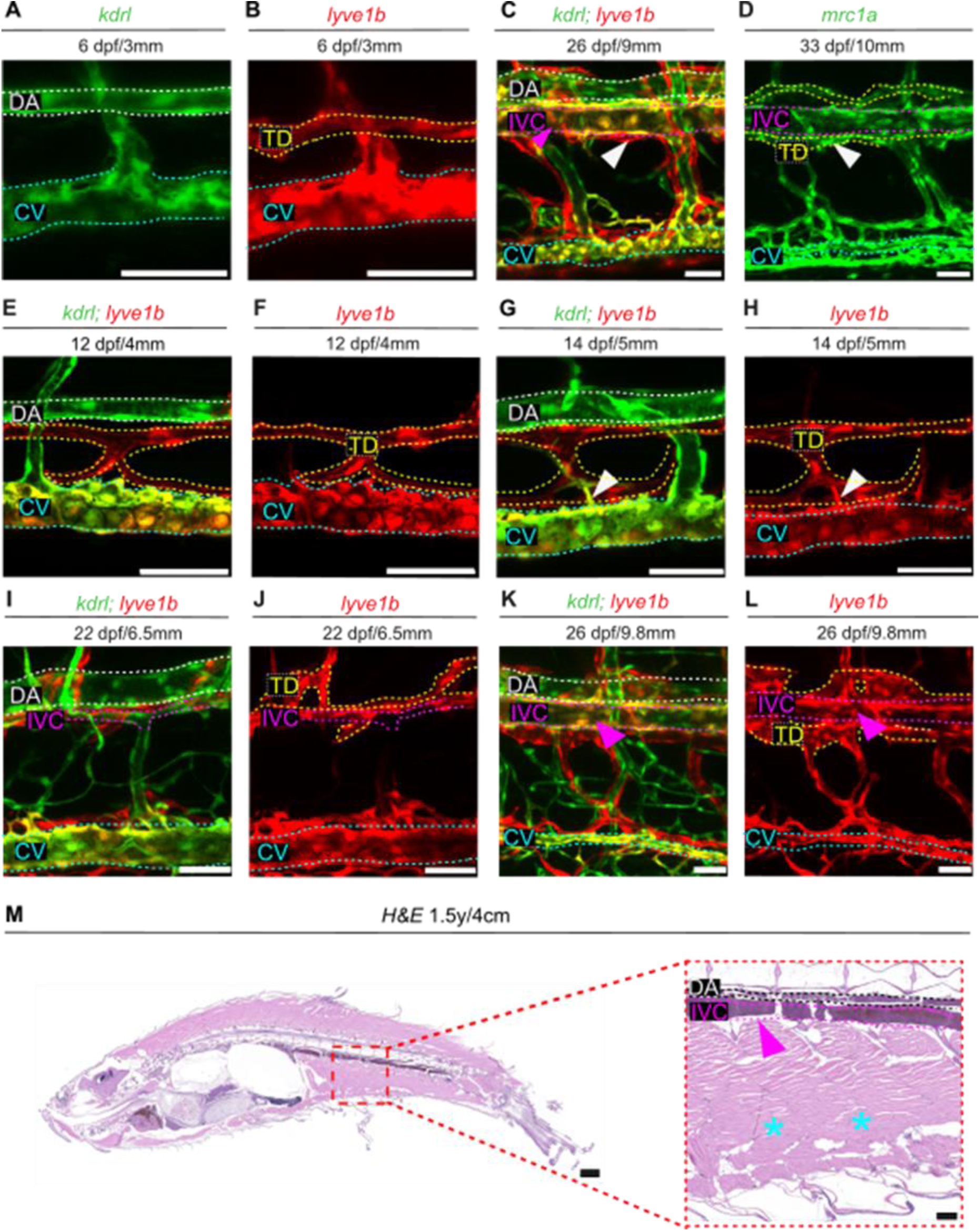
Progressive transition from the CV to the IVC during zebrafish metamorphosis. (**A,B**) At 6 dpf/3mm Tg(kdrl:GFP) labels the DA (white dashed lines) and CV (light-blue dashed lines), while Tg(lyve1b:dsRed) marks the TD (yellow dashed lines) and CV. (C) At 26 dpf/9mm Tg(kdrl:GFP; lyve1b:dsRed), reveals the emergence of the IVC (magenta dashed lines, magenta arrowhead) partially wrapped by lymphatic vessels (white arrowhead). (D) Tg(mrc1a:GFP) fish at 33 dpf/10mm shows labeling of the IVC (magenta dashed lines), TD (yellow dashed lines), and CV (light blue dashed lines). (E,F) At 12 dpf/4mm, Tg(kdrl:GFP;lyve1b:dsRed) larvae show kdrl/lyve1b colocalization in the CV, kdrl+ labeling of the DA, and lyve1b labeling of lymphatic and CV ECs. (G,H) At 14 dpf/5mm kdrl:GFP+;lyve1b:dsRed+ sprouts emerge from the CV (white arrowhead). (I-L) At 22 dpf/6.5mm (I,J) and 26 dpf/9.8mm (K,L), the forming IVC is highlighted by kdrl:GFP and lyve1b:dsRed (magenta dashed lines, magenta arrowhead). (M) Sagittal H&E-stained section of a 1.5-year-old fish showing the IVC (magenta dashed line) while the CV is no longer detected (light-blue asterisks). Scale bars, 25 µm (A,B), 50 µm (C,D,E-L,M), Cardinal vein (CV), Dorsal Aorta (DA), white dashed lines; Thoracic duct (TD), yellow dashed lines; Inferior Vena Cava (IVC), magenta dashed lines.

**Fig. S2.**
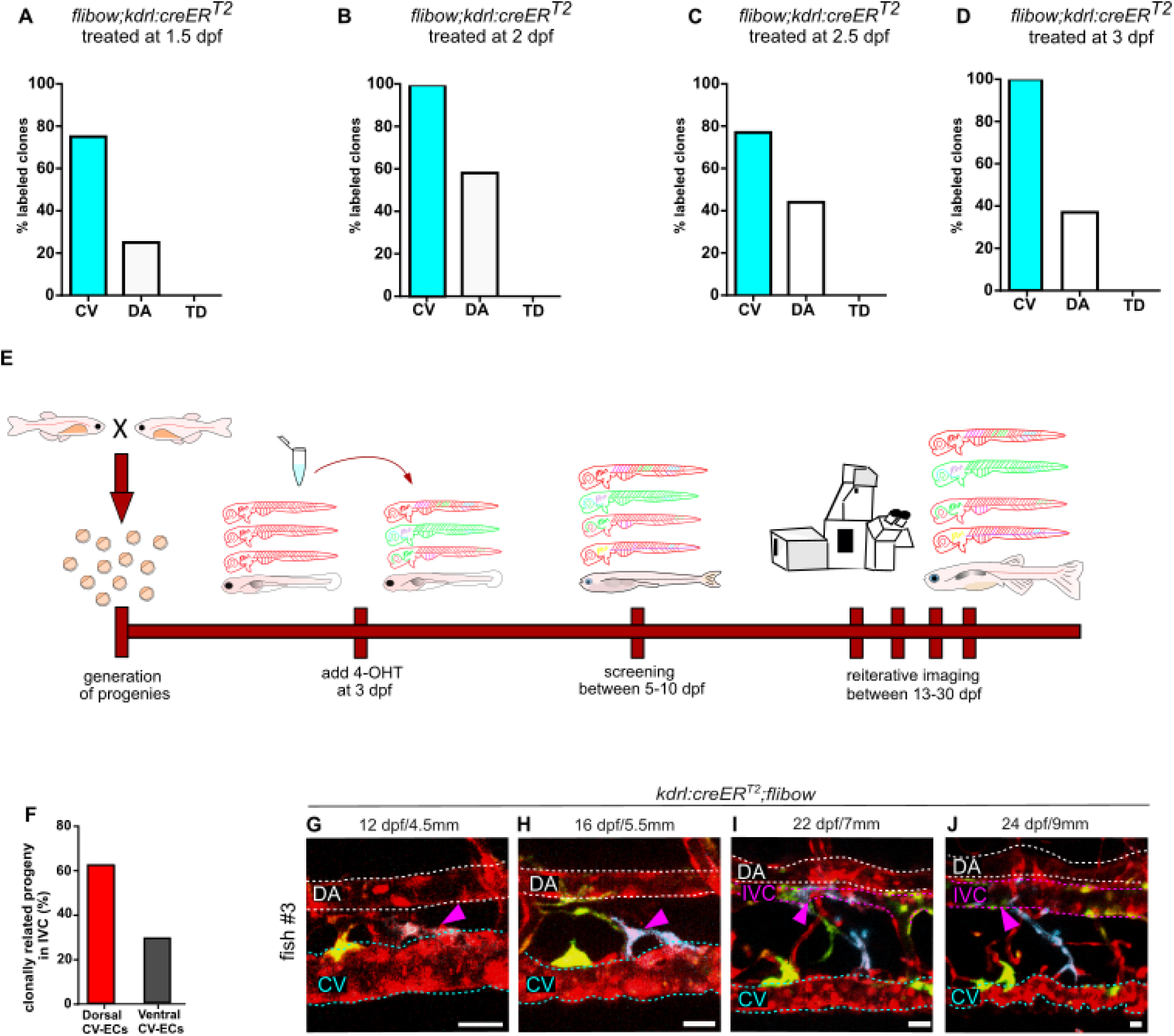
Lineage tracing reveals progressive contribution of CV-ECs to the IVC. (**A-D**) Quantification of labeled EC clones in Tg(kdrl:CreER^T2^;flibow) fish treated with 4-OH at 1.5, 2, 2.5, and 3dpf. (E) Schematic representation of the flibow-based lineage tracing approach used in this study. (F) Proportion of IVC clones derived from dorsal vs. ventral CV-ECs. (G-J) Reiterative imaging of the same Tg(kdrl:CreER^T2^;flibow) fish. (G) A labeled CV-EC clone is detected at CV at 12 dpf/4.5mm (magenta arrowhead), in CV sprouts at 16 dpf/5.5mm (H), in the nascent IVC at 22 dpf/7mm (I), and in the mature IVC at 24 dpf/9mm (J). Scale bars 50 µm (G-J). Cardinal vein (CV), light-blue dashed lines; Dorsal Aorta (DA), white dashed lines; Inferior Vena Cava (IVC), magenta dashed lines.

**Fig. S3.**
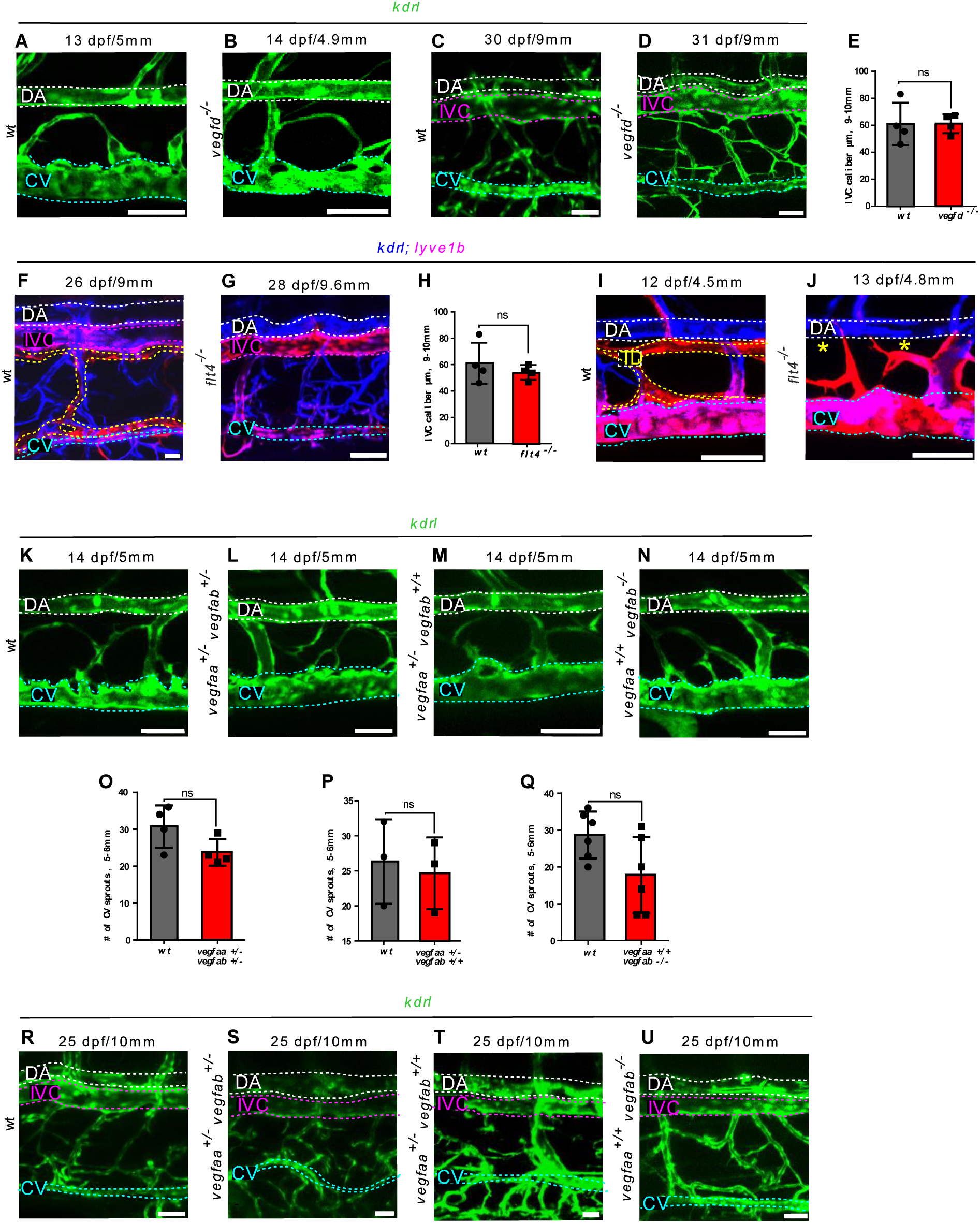
Assessment of Vegf pathway components in CV sprouting and IVC formation. (**A-B**) Tg(kdrl:EGFP); vegfd ^-/-^ and wt siblings show comparable CV sprouting. (C-D) IVC formation is unaffected in vegfd mutants. IVC caliber is quantified in (E), showing no difference between wt and vegfd ^-/-^ (n=5, t-test, p > 0,05). (F-G). IVC forms normally in Tg(kdrl:BFP;lyve1b:dsRed);flt4^-/-^ animals; quantified in (H) (n=4, t-test, p > 0,05). (I-J) CV sprouts are present in Tg(kdrl:BFP;lyve1b:dsRed);flt4^-/-^, though lymphatic vessels are missing (yellow asterisks), confirming loss of flt4 function. (K–N) CV sprouting at 14 dpf/5mm is similar across Tg(kdrl:EGFP) animals with vegfaa⁺/⁺;vegfab⁺/⁺, vegfaa⁺/⁻;vegfab⁺/⁻, vegfaa⁺/⁻;vegfab⁺/⁺, and vegfaa⁺/⁺; vegfab⁻/⁻ genotypes; quantified in (O-Q) (n=4; n=3; n=6; t-test, p>0,05). (R-U) IVC is normally present at 25 dpf/10mm in vegfaa^+/+^;vegfab^+/+^, vegfaa^+/-^;vegfab^+/-^, vegfaa^+/-^;vegfab^+/+^ and vegfaa^+/+^;vegfab^-/-^ animals. Scale bars 50 µm (A-D, K-N, R-U). Cardinal vein (CV), light-blue dashed lines; Dorsal Aorta (DA), white dashed lines; Inferior Vena Cava (IVC), magenta dashed lines.

**Movie S1.**

Time-lapse video using Light sheet microscopy of Tg(fli1:EGFP; gata1:dsRed) at 18 dpf/5.5mm trunk showing the blood flow direction. Erythrocytes in the CV (white arrowheads) move toward the heart, while in the DA, erythrocytes (magenta arrowheads) flow caudally toward the tail fin. The video spans 1000 frames on a single Z-stack. Cardinal vein (CV); Dorsal Aorta (DA); Inferior Vena Cava (IVC).

**Movie S2.**

Time-lapse video using Light sheet microscopy of Tg(fli1:EGFP; gata1:dsRed) at 25 dpf/9mm showing the presence of erythrocytes in the IVC (magenta arrowheads) running towards the heart, indicating a venous identity. In the DA, erythrocytes (white arrowheads) flow towards the tail fin. The video spans 1000 frames on a single Z-stack. Cardinal vein (CV); Dorsal Aorta (DA); Inferior Vena Cava (IVC).

**Movie S3.**

Time-lapse video using confocal microscopy of Tg(kdrl:BFP;lyve1b:dsRed;runx1:GFP) trunk area at 4 dpf, showing the DA (kdrl+), CV (kdrl+/lyve1+) and runx1+ cells (white arrowheads) detected within EC pockets in the CV-CHT, and in circulation (yellow arrowhead). The video is played at 1.8 fps. Cardinal vein (CV); Dorsal Aorta (DA); Inferior Vena Cava (IVC).

**Movie S4.**

Time-lapse video of Tg(kdrl:BFP;lyve1b:dsRed;runx1:GFP) trunk area using confocal microscopy of 25 dpf/9mm animals. runx1+ cells are detected only in circulation (white arrowheads), indicating that the IVC does not harbor HSCs. The video is played at 1.8 fps. Cardinal vein (CV); Dorsal Aorta (DA); Inferior Vena Cava (IVC).

## Materials and Methods

### Zebrafish husbandry and strains

Zebrafish were raised by standard methods and handled according to the Weizmann Institute Animal Care and Use Committee guidelines. Zebrafish lines used in this study were *Tg(fli1:EGFP)^y1^*(*55*), *Tg(lyve1b:dsRed)^nz101^*(*56*)*, Tg(kdrl:EGFP)^s843^*(*57*)*, Tg(kdrl:tagBFP)^mu293^*(*57*)*, Tg(mrc1a:EGFP)^y251^*(*25*) *Tg(gata1a:dsRed)^sd2^*(*58*)*,, Tg(flibow)*(*24*)*, Tg(kdrl:cre-ER^T2^)^fb13^*(*26*)*, Tg(hsp70l:creER^T2^)*(*24*)*, Tg(vegfaa^bn1^; vegfab^bns92^)*(*33*)*, Tg(flt4^um203^)*(*59*)*, Tg(vegfd^bns257^)*(*54*)*,, TgBAC(flt1:YFP)^hu4624^* (*60*)*, Tg(vegfaa:EGFP)*(*61*) *Tg(dll4:gal4)^mut106^*(*40*)*, Tg(UAS:Kaede)*(*62*) *Tg(pdgfrb:Gal4)*(*63*) *Tg(pdgfrb:EGFP)*(*63*)*, Tg(uas:NTR_cherry)*(*64*) *Tg(runx1:GFP)*(*46*) were previously described. For imaging, we used either a *casper (roy^−/−^; nacre^−/−^)*(*65*) background, or embryos were treated for 5 days with 0.003% N-phenylthiourea (PTU) (Sigma, St Louis, MO) to inhibit melanin pigment formation.

### Lineage Tracing Analyses

For lineage tracing experiments, we used the previously generated *Tg(flibow)* line (*24*). To achieve efficient labeling of the CV and DA, Cre expression was induced at 3 dpf. For *Tg(kdrl:creER^T2^)* mediated Cre expression, embryos were treated with 5µM 4-hydroxytamoxifen for 24 hours. For *Tg(hsp70l:creER^T2^)* induction, embryos were pre-heated at 36°C for 15 mins to activate the *hsp70l* promoter, followed by a 45-minute combined heat (36°C) and 5µM 4-hydroxytamoxifen (diluted in E3 medium) treatment. After treatment, embryos were rinsed three times in fresh E3 medium and returned to system water. Animals with labeled cells were screened at 5-10dpf, and selected individuals were imaged at the appropriate IVC developmental stages and maintained in fresh water until the experimental endpoint.

*Tg(kdrl:creER^T2^; flibow*) fish were treated with 4-hydroxytamoxifen (4-OHT) at 1.5, 2, 2.5, and 3 dpf and analyzed at the first time point (10-14 dpf) to assess the distribution of labeled clones in different vessel types. At 1.5 dpf among 5 fish, 4 showed labeling in venous ECs, 2 in arterial ECs and none in lymphatic ECs. At 2dpf, of 13 fish analyzed, 12 exhibited venous labeling, 7 exhibited arterial labeling, and 0 exhibited lymphatic labeling. At 2.5 dpf, among 9 fish, 7 showed venous labeling, 4 arterial, and none lymphatic. At 3dpf, all 32 fish showed venous labeling, 12 had arterial labeling, and none exhibited lymphatic labeling.

To calculate the number of ventral and dorsal ECs contributing to IVC formation, fish were analyzed at the first time point (10-14 dpf) and scored for the location of the labeled ECs. Of the labeled cells, 33 dorsal cells give rise to IVC, whereas only 8 ventral ECs were found to contribute to the IVC.

### Microinjections and uptake assays

Microangiography was performed on anesthetized juvenile animals using microinjection glass capillaries, as previously described (*66*). The following compounds were injected: Qtracker705 (Invitrogen, Q21061MP), Tetramethylrhodamine Dextran 10k (Thermofisher, D7139), Fluorescein isothiocyanate (FITC) Dextran (500,000 MW) (Sigma, 46947) and Transferrin (Invitrogen, T13342) The injected animals were allowed to recover in fresh water for 5 mins, following which they were anesthetized and processed for confocal microscopy.

### Histology sample preparation

Samples from 8mm/∼20 dpf and 1.5-year-old zebrafish were fixed in 4% paraformaldehyde (PFA) overnight at 4°C. Tissue processing was performed using a Leica ASP300S processor with an automatic protocol. The dehydration process began with 70% ethanol for 45 minutes, followed by three steps of 96% ethanol for 30 minutes each, and two 100% ethanol for 30 minutes each. Subsequently, the samples were washed with ethanol and Medite X-tra-solv xylene substitute for 45 minutes, followed by two additional washes with Medite X-tra-solv for 60 minutes each. Finally, the wax was changed three times at 57°C for 60 minutes each before embedding the samples in Leica Paraplast paraffin.

### H&E Staining Protocol

H&E staining was performed using an automatic stainer (Giotto, Diapath). Samples were initially cleared using Medite X-tra-solve xylene substitute in three 5-minute washes, followed by rehydration, with sequential 5-minute incubations in 100%, 96%, and 70% ethanol. After brief rinses in water and distilled water (1 minute each) slides were stained with Hematoxylin for 5 minutes, followed by a sequence of washes: Tween (2 minutes), acidified 70% ethanol (10 seconds), two tween washes (3 minutes each), distilled water (1 minute), and 70% ethanol (3 minutes). Slides were then stained with eosin. Dehydration was completed with two 2-minute washes in 96% ethanol and two 3-minute washes in 100% ethanol. The protocol concluded with two 3-minute clearing steps using Medite X-tra-solv xylene substitute.

### Confocal microscopy

Confocal imaging was performed using a Zeiss LSM880 upright confocal microscope equipped with a water-immersed W-Plan Apochromat 20x NA 1.0 objective lens. Euthanized animals were mounted in 1.5% w/v low-melting-point agarose. For reiterative imaging a custom-built imaging chamber was utilized(*24*). Z-stacks were acquired at 2-3 μm intervals. Larger fields of view were obtained via tile-scanning, and images were stitched using Imaris Stitcher 9.3 or Zeiss Zen software. *Tg*(*flibow)* confocal imaging and lambda stack acquisition were performed as previously described (*24*).

### Time-lapse Imaging

Time-lapse recordings of blood flow were acquired using a Light Sheet Z.1 microscope (Zeiss Ltd.) equipped with a 10X/0.1 lens and dual PCO-Edge sCMOS cameras. Larvae were anesthetized, embedded in 1.5% low-melting-point agarose, and mounted in glass capillaries. Subsequently, animals were suspended in egg water using the plunger. Images were captured over 1000 frames within a single Z-stack. Analysis was conducted using Zen Blue and ARIVIS software.

### Image processing

Confocal images were processed off-line using Fiji version of ImageJ (NIH). The images shown in this study are single views, 2D reconstructions of collected z-series stacks. Co-localization thresholds were set manually. *Tg(flibow)* images were processed using Imaris 9.3(Bitplane). Each of the 11 channels was given RGB values corresponding to the wavelength collected. We chose six channels, leaving out those that were redundant or displayed a poor signal/noise ratio, as described previously(*24*). The H&E images were processed using CaseViewer.

### Statistical Analyses

Comparison of two samples was done using unpaired two-tailed Student’s t-test assuming equal variances from at least three independent experiments unless stated otherwise. Statistical significance for three or more samples was calculated via one-way ANOVA, followed by Tukey’s or Dunnett’s multiple comparisons test, unless stated otherwise. All data are reported as mean values ± SEM and were analyzed using Prism 6 software (GraphPad Software, Incorporated, La Jolla, CA, USA).

