## Supplementary material for "A Conserved Venous Remodeling Program Governs Inferior Vena Cava Formation in Zebrafish": Table_1

| mple num | Experiment folder | Sample name | Exp date | Induction conditions | CRE line | Lymphatic | vISV | CV | DA | IVC | OBSERVATION |
|---|---|---|---|---|---|---|---|---|---|---|---|
| 1 | 2024-4_5_flibowkdrlcreert_treatedat | f1_12dpf_midtrunkblucell_2024-05-17T19-01-21.023 | | treated at 3dpf 24h tamox | kdrl | na | x | x | na | \ | |
| 2 | 2024-4_5_flibowkdrlcreert_treatedat | f1_16dpf_midtrunkblucel_5mml_2024-05-17T19-04-02.895 | | treated at 3dpf 24h tamox | kdrl | na | x | x | na | \ | |
| 3 | 2024-4_5_flibowkdrlcreert_treatedat | f1_22dpf_midtrunkblucel_8mml_2024-05-17T19-08-37.872 | | treated at 3dpf 24h tamox | kdrl | na | x | x | na | x | |
| 3 | 2024-4_5_flibowkdrlcreert_treatedat | f1_22dpf_midtrunkblucel_8mml_2024-05-17T19-08-37.872 | | treated at 3dpf 24h tamox | kdrl | na | x | x | na | x | |
| 4 | 2024-4_5_flibowkdrlcreert_treatedat | f2_12dpf_analpore_2024-07-28T18-01-57.979 | | treated at 3dpf 24h tamox | kdrl | na | na | x | na | \ | |
| 7 | 2024-4_5_flibowkdrlcreert_treatedat | f2_16dpf_analpore.5mmczi_2024-07-28T18-16-46.688 | | treated at 3dpf 24h tamox | kdrl | na | x | x | na | \ | |
| 11 | 2024-4_5_flibowkdrlcreert_treatedat | f2_22dpf_analpore.6mmczi_2024-07-28T18-23-37.545 | | treated at 3dpf 24h tamox | kdrl | na | x | x | na | \ | |
| 15 | 2024-4_5_flibowkdrlcreert_treatedat | f2_24dpf_analpore.7mmczi_2024-07-28T18-34-29.172 | | treated at 3dpf 24h tamox | kdrl | na | x | x | na | x | |
| 19 | 2024-4_5_flibowkdrlcreert_treatedat | f2_33dpf_analpore.8.5mmczi_2024-07-28T18-52-08.183 | | treated at 3dpf 24h tamox | kdrl | na | na | na | na | na | |
| 5 | 2024-4_5_flibowkdrlcreert_treatedat | f2_12dpf_midtrunk_2024-07-28T18-07-11.717 | | treated at 3dpf 24h tamox | kdrl | na | x | x | na | \ | |
| 8 | 2024-4_5_flibowkdrlcreert_treatedat | f2_16dpf_midtrunk5mm_2024-07-28T18-18-07.736 | | treated at 3dpf 24h tamox | kdrl | na | x | x | na | \ | |
| 12 | 2024-4_5_flibowkdrlcreert_treatedat | f2_22dpf_midtrunk6mm_2024-07-28T18-27-33.731 | | treated at 3dpf 24h tamox | kdrl | na | na | x | na | \ | |
| 16 | 2024-4_5_flibowkdrlcreert_treatedat | f2_24dpf_midtrunk7.7mm_2024-07-28T18-47-57.822 | | treated at 3dpf 24h tamox | kdrl | na | x | x | na | \ | |
| 6 | 2024-4_5_flibowkdrlcreert_treatedat | f2_12dpf_tail_2024-07-28T18-14-52.864 | | treated at 3dpf 24h tamox | kdrl | na | na | x | na | \ | |
| 10 | 2024-4_5_flibowkdrlcreert_treatedat | f2_16dpf_tail5mm_2024-07-28T18-21-44.465 | | treated at 3dpf 24h tamox | kdrl | na | na | x | x | \ | |
| 14 | 2024-4_5_flibowkdrlcreert_treatedat | f2_22dpf_tail6mm_2024-07-28T18-30-51.603 | | treated at 3dpf 24h tamox | kdrl | na | na | x | na | \ | |
| 18 | 2024-4_5_flibowkdrlcreert_treatedat | f2_24dpf_tail7mm_2024-07-28T18-50-14.807 | | treated at 3dpf 24h tamox | kdrl | na | na | x | na | \ | |
| 9 | 2024-4_5_flibowkdrlcreert_treatedat | f2_16dpf_postanalpore.5mmczi_2024-07-28T18-19-48.897 | | treated at 3dpf 24h tamox | kdrl | na | na | x | na | \ | |
| 13 | 2024-4_5_flibowkdrlcreert_treatedat | f2_22dpf_postanalpore.7mmczi_2024-07-28T18-29-27.331 | | treated at 3dpf 24h tamox | kdrl | na | x | x | na | \ | |
| 17 | 2024-4_5_flibowkdrlcreert_treatedat | f2_24dpf_postanalpore.7.5mmczi_2024-07-28T18-49-14.0 | | treated at 3dpf 24h tamox | kdrl | na | x | x | na | x | |
| 24 | 2024-4_5_flibowkdrlcreert_treatedat | f3_16dpf_midtrunk1cell_5mm_2024-07-28T18-59-02.905 | | treated at 3dpf 24h tamox | kdrl | na | na | x | na | \ | |
| 25 | 2024-4_5_flibowkdrlcreert_treatedat | f3_22dpf_midtrunk1cell_8mm_2024-07-28T19-00-23.672 | | treated at 3dpf 24h tamox | kdrl | na | x | x | na | na | |
| 26 | 2024_5_7flibowkdrlcreerttreatedat3dp | f1_13dpf_2024-07-28T16-33-01.589 | | treated at 3dpf 24h tamox | kdrl | na | na | x | na | \ | |
| 27 | 2024_5_7flibowkdrlcreerttreatedat3dp | f1_22dpf_2024-07-28T16-37-28.174 | | treated at 3dpf 24h tamox | kdrl | na | x | na | na | \ | |
| 28 | 2024_5_7flibowkdrlcreerttreatedat3dp | f1_22dpf_midtrunk_2024-07-28T16-39-27.487 | | treated at 3dpf 24h tamox | kdrl | na | x | x | na | \ | |
| 29 | 2024_5_7flibowkdrlcreerttreatedat3dp | f2_13dpf_2024-07-28T16-41-10.656 | | treated at 3dpf 24h tamox | kdrl | na | na | x | na | \ | |
| 30 | 2024_5_7flibowkdrlcreerttreatedat3dp | f2_22dpf_2024-07-28T16-44-10.817 | | treated at 3dpf 24h tamox | kdrl | na | na | x | na | \ | |
| 31 | 2024_5_7flibowkdrlcreerttreatedat3dp | f4_13dpf_2024-07-28T16-48-29.890 | | treated at 3dpf 24h tamox | kdrl | na | x | x | x | \ | |
| 32 | 2024_5_7flibowkdrlcreerttreatedat3dp | f4_22dpf_2024-07-28T16-51-13.202 | | treated at 3dpf 24h tamox | kdrl | na | x | x | x | \ | |
| 33 | | | | | | | | | | | |
| 34 | 2024_5_7flibowkdrlcreerttreatedat3dp | f5_13dpf_2024-07-28T16-53-10.178 | | treated at 3dpf 24h tamox | kdrl | na | x | x | na | \ | |
| 35 | 2024_5_7flibowkdrlcreerttreatedat3dp | f6_13dpf_dagood_4mm_2024-05-17T16-04-41.826 | | treated at 3dpf 24h tamox | kdrl | na | na | na | x | \ | |
| 36 | 2024_5_7flibowkdrlcreerttreatedat3dp | f6_22dpf_dagood_5mm_2024-05-17T16-10-31.220 | | treated at 3dpf 24h tamox | kdrl | na | na | na | x | \ | |
| 37 | 2024_5_7flibowkdrlcreerttreatedat3dp | f6_25dpf_DAgood_6mm_2024-05-22T16-17-07.036 | | treated at 3dpf 24h tamox | kdrl | na | na | na | x | na | |
| 38 | 2024_5_7flibowkdrlcreerttreatedat3dp | f6_30dpf_DAgood_8.1mm_zoomczi_2024-05-25T20-05-01 | | treated at 3dpf 24h tamox | kdrl | na | na | na | x | na | |
| 39 | 2024_5_7flibowkdrlcreerttreatedat3dp | f7_13dpf_2cellanalpore_2024-07-28T16-55-23.972 | | treated at 3dpf 24h tamox | kdrl | na | na | x | na | \ | |
| 41 | 2024_5_7flibowkdrlcreerttreatedat3dp | f7_22dpf_2cellanalpore_2024-07-28T17-01-04.436 | | treated at 3dpf 24h tamox | kdrl | na | x | x | na | \ | |
| 40 | 2024_5_7flibowkdrlcreerttreatedat3dp | f7_13dpf_midtrunk_2024-07-28T16-58-24.396 | | treated at 3dpf 24h tamox | kdrl | na | x | x | na | \ | |
| 42 | 2024_5_7flibowkdrlcreerttreatedat3dp | f7_22dpf_midtrunk_2024-07-28T17-02-37.660 | | treated at 3dpf 24h tamox | kdrl | na | x | x | na | \ | |
| 43 | 2024_5_7flibowkdrlcreerttreatedat3dp | f8_13dpf_analopore_2024-07-28T17-04-13.709 | | treated at 3dpf 24h tamox | kdrl | na | x | x | na | \ | |
| 45 | 2024_5_7flibowkdrlcreerttreatedat3dp | f8_22dpf_analopore_2024-07-28T17-09-32.510 | | treated at 3dpf 24h tamox | kdrl | na | x | x | na | \ | |
| 44 | 2024_5_7flibowkdrlcreerttreatedat3dp | f8_13dpf_midtrunk_2024-07-28T17-08-27.638 | | treated at 3dpf 24h tamox | kdrl | na | x | x | na | \ | |
| 46 | 2024_5_7flibowkdrlcreerttreatedat3dp | f9_13dpf_2024-07-28T17-11-56.182 | | treated at 3dpf 24h tamox | kdrl | na | x | x | na | \ | |
| 47 | 2024_5_7flibowkdrlcreerttreatedat3dp | f9_22dpf_2024-07-28T17-14-45.792 | | treated at 3dpf 24h tamox | kdrl | na | x | x | na | \ | |
| 48 | 2024_5_7flibowkdrlcreerttreatedat3dp | f10_14dpf_DAimport_2024-05-17T16-42-59.580 | | treated at 3dpf 24h tamox | kdrl | na | na | na | x | \ | |
| 49 | 2024_5_7flibowkdrlcreerttreatedat3dp | f10_22dpf_DAimport_2024-05-17T16-24-17.614 | | treated at 3dpf 24h tamox | kdrl | na | na | na | x | \ | |
| 50 | 2024_5_7flibowkdrlcreerttreatedat3dp | f10_25dpf_daimport_zoom_1_2024-05-22T16-41-44.127 | | treated at 3dpf 24h tamox | kdrl | na | na | na | x | na | |
| 51 | 2024_5_7flibowkdrlcreerttreatedat3dp | f10_30dpf_daimport_zoom_8.5mm_2024-05-25T19-50-11 | | treated at 3dpf 24h tamox | kdrl | na | na | na | x | na | |
| 53 | 2022.10.27flibowkdrlcreert treatedat2.5dpf10uM | | | | | | | | | | |
| 54 | 2022.10.27flibowkdrlcreert treatedat2 | f1 flibow kdrlcreert treated at 2.5dpf 10um 8dpf | | treated at 2.5dpf 24h tamox | kdrl | na | x | x | na | \ | |
| 55 | 2022.10.27flibowkdrlcreert treatedat2 | f1 flibow kdrlcreert treated at 2.5dpf 10um 14dpf | | treated at 2.5dpf 24h tamox | kdrl | na | x | x | na | \ | |
| 56 | 2022.10.27flibowkdrlcreert treatedat2 | f1 flibow kdrlcreert treated at 2.5dpf 10um 23dpf 9mm | | treated at 2.5dpf 24h tamox | kdrl | na | x | x | na | na | |
| 57 | 2022.10.27flibowkdrlcreert treatedat2 | f2 flibow kdrlcreert treated at 2.5dpf 10um 8dpf | | treated at 2.5dpf 24h tamox | kdrl | na | x | x | na | \ | |
| 58 | 2022.10.27flibowkdrlcreert treatedat2 | f2 flibow kdrlcreert treated at 2.5dpf 10um 23dpf10mm sar | | treated at 2.5dpf 24h tamox | kdrl | na | x | x | na | x | |
| 60 | 2022.10.27flibowkdrlcreert treatedat2 | f3 flibow kdrlcreert treated at 2.5dpf 10um 12dpf cell anal | | treated at 2.5dpf 24h tamox | kdrl | na | x | na | x | \ | |
| 62 | 2022.10.27flibowkdrlcreert treatedat2 | f3 flibow kdrlcreert treated at 2.5dpf 10um 23dpf cell anal | | treated at 2.5dpf 24h tamox | kdrl | na | x | na | x | x | |
| 59 | 2022.10.27flibowkdrlcreert treatedat2 | f3 flibow kdrlcreert treated at 2.5dpf 10um 12dpf 3cells mid | | treated at 2.5dpf 24h tamox | kdrl | na | na | x | na | \ | |
| 61 | 2022.10.27flibowkdrlcreert treatedat2 | f3 flibow kdrlcreert treated at 2.5dpf 10um 23dpf 2 cells mi | | treated at 2.5dpf 24h tamox | kdrl | na | x | x | na | na | |
| 63 | 2022.10.27flibowkdrlcreert treatedat2 | f4 flibow kdrlcreert treated at 2.5dpf 10um 12dpf 2 cells mi | | treated at 2.5dpf 24h tamox | kdrl | na | na | x | na | \ | |
| 66 | 2022.10.27flibowkdrlcreert treatedat2 | f4 flibow kdrlcreert treated at 2.5dpf 10um 14dpf 2 cells mi | | treated at 2.5dpf 24h tamox | kdrl | na | na | x | na | x | |
| 69 | 2022.10.27flibowkdrlcreert treatedat2 | f4 flibow kdrlcreert treated at 2.5dpf 10um 23dpf 1 cells mi | | treated at 2.5dpf 24h tamox | kdrl | na | x | na | na | x | |
| 64 | 2022.10.27flibowkdrlcreert treatedat2 | f4 flibow kdrlcreert treated at 2.5dpf 10um 12dpf cell close | | treated at 2.5dpf 24h tamox | kdrl | na | na | x | na | \ | |
| 67 | 2022.10.27flibowkdrlcreert treatedat2 | f4 flibow kdrlcreert treated at 2.5dpf 10um 14dpf cell close | | treated at 2.5dpf 24h tamox | kdrl | na | na | x | na | \ | |
| 72 | 2022.10.27flibowkdrlcreert treatedat2 | f4 flibow kdrlcreert treated at 2.5dpf 10um 23dpf cell close | | treated at 2.5dpf 24h tamox | kdrl | na | na | na | na | x | |
| 65 | 2022.10.27flibowkdrlcreert treatedat2 | f4 flibow kdrlcreert treated at 2.5dpf 10um 12dpf da and pc | | treated at 2.5dpf 24h tamox | kdrl | na | na | x | x | \ | |
| 68 | 2022.10.27flibowkdrlcreert treatedat2 | f4 flibow kdrlcreert treated at 2.5dpf 10um 14dpf da and pc | | treated at 2.5dpf 24h tamox | kdrl | na | na | x | x | x | |
| 73 | 2022.10.27flibowkdrlcreert treatedat2 | f5 flibow kdrlcreert treated at 2.5dpf 10um 12dpf all trunk | | treated at 2.5dpf 24h tamox | kdrl | na | x | x | na | \ | |
| 74 | 2022.10.27flibowkdrlcreert treatedat2 | f5 flibow kdrlcreert treated at 2.5dpf 10um 12dpf all trunk | | treated at 2.5dpf 24h tamox | kdrl | na | x | x | x | \ | |
| 75 | 2022.10.27flibowkdrlcreert treatedat2 | f5 flibow kdrlcreert treated at 2.5dpf 10um 12dpf all trunk | | treated at 2.5dpf 24h tamox | kdrl | na | x | x | na | \ | |
| 76 | 2022.10.27flibowkdrlcreert treatedat2 | f6 flibow kdrlcreert treated at 2.5dpf 10um 12dpf all trunk | | treated at 2.5dpf 24h tamox | kdrl | na | na | x | x | \ | |
| 77 | 2022.10.27flibowkdrlcreert treatedat2 | f6 flibow kdrlcreert treated at 2.5dpf 10um 12dpf all trunk | | treated at 2.5dpf 24h tamox | kdrl | na | na | x | x | \ | |
| 78 | 2022.10.27flibowkdrlcreert treatedat2 | f6 flibow kdrlcreert treated at 2.5dpf 10um 12dpf all trunk | | treated at 2.5dpf 24h tamox | kdrl | na | na | x | x | \ | |
| 80 | 2022.10.27flibowkdrlcreert treatedat2 | f6 flibow kdrlcreert treated at 2.5dpf 10um 23dpf all trunk | | treated at 2.5dpf 24h tamox | kdrl | na | na | x | x | x | |
| 81 | 2022.10.27flibowkdrlcreert treatedat2 | f6 flibow kdrlcreert treated at 2.5dpf 10um 23dpf all trunk | | treated at 2.5dpf 24h tamox | kdrl | na | na | x | x | x | |
| 82 | 2022.10.27flibowkdrlcreert treatedat2 | f6 flibow kdrlcreert treated at 2.5dpf 10um 23dpf all trunk | | treated at 2.5dpf 24h tamox | kdrl | na | na | x | x | x | |
| 79 | 2022.10.27flibowkdrlcreert treatedat2 | f6 flibow kdrlcreert treated at 2.5dpf 10um 12dpfblu anal p | | treated at 2.5dpf 24h tamox | kdrl | na | x | x | na | \ | |
| 83 | 2022.10.27flibowkdrlcreert treatedat2 | f6 flibow kdrlcreert treated at 2.5dpf 10um 23dpfblu anal p | | treated at 2.5dpf 24h tamox | kdrl | na | na | na | na | x | |
| 84 | 2024_3_21flibowkdrlcreert treated at 3dpf | | | | | | | | | | |
| 85 | 2024_3_21flibowkdrlcreert treated at | f10_tail_11dpf_2024-05-21T15-54-53.750 | | treated at 3dpf | kdrl | na | na | x | na | \ | |
| 86 | 2024_3_21flibowkdrlcreert treated at | f10_tail_29dpf_9mm_2024-05-21T15-56-24.648 | | treated at 3dpf | kdrl | na | na | na | na | x | |
| 87 | 2024_3_21flibowkdrlcreert treated at | f9_analporelgood_11dpf_2024-05-21T15-03-18.228 | | treated at 3dpf | kdrl | na | x | x | na | \ | |
| 88 | 2024_3_21flibowkdrlcreert treated at | f9_analporelgood_14dpf_5mm_2024-05-21T15-13-25.237 | | treated at 3dpf | kdrl | na | x | x | na | \ | |
| 89 | 2024_3_21flibowkdrlcreert treated at | f9_analporelgood_21dpf_7.5mm_2024-05-21T15-08-41.14 | | treated at 3dpf | kdrl | na | x | x | na | x | |
| 90 | 2024_3_21flibowkdrlcreert treated at 3dpf | | | | | | | | | | |
| 91 | | | | | | | | | | | |
| 92 | 2024_3_21flibowkdrlcreert treated at | f10_midtrunk_11dpf_2024-05-21T15-36-15.337 | | treated at 3dpf | kdrl | na | x | x | na | \ | |
| 94 | 2024_3_21flibowkdrlcreert treated at | f10_midtrunk_14dpf_5mm_2024-05-21T15-38-09.985 | | treated at 3dpf | kdrl | na | x | x | na | \ | |
| 95 | 2024_3_21flibowkdrlcreert treated at | f10_midtrunk_21dpf_5.5mm_2024-05-21T15-39-37.082 | | treated at 3dpf | kdrl | na | x | x | na | \ | |
| 96 | 2024_3_21flibowkdrlcreert treated at | f10_midtrunk_29dpf_9mm_2024-05-21T15-41-38.586 | | treated at 3dpf | kdrl | na | x | x | na | x | |
| 97 | 2024_3_21flibowkdrlcreert treated at | f12_midtrunksuper_12dpf | | treated at 3dpf | kdrl | na | na | x | na | \ | |
| 98 | 2024_3_21flibowkdrlcreert treated at | f12_midtrunksuper_14dpf_5mm | | treated at 3dpf | kdrl | na | na | x | na | \ | |
| 99 | 2024_3_21flibowkdrlcreert treated at | f12_midtrunksuper2_21dpf_6.2mm | | treated at 3dpf | kdrl | na | na | x | na | \ | |
| 100 | 2024_3_21flibowkdrlcreert treated at | f12_midtrunksuper2_29dpf_9mm | | treated at 3dpf | kdrl | na | na | x | na | x | |
| 101 | 2021.2.21 flibow kdrlcreert treated at 1.5dpf tamoxifen 10um | | | | | | | | | | |
| 103 | 2021.2.21 flibow kdrlcreert treated at | f1 12dpf_treated at 1.5dpf full trunk pcv cells | | treated at 1.5dpf | kdrl | na | x | x | na | \ | |
| 104 | 2021.2.21 flibow kdrlcreert treated at | f1 12dpf_treated at 1.5dpf full trunk pcv cells | | treated at 1.5dpf | kdrl | na | x | x | na | \ | |
| 105 | 2021.2.21 flibow kdrlcreert treated at | f1 12dpf_treated at 1.5dpf full trunk pcv cells | | treated at 1.5dpf | kdrl | na | x | x | na | \ | |
| 106 | 2021.2.21 flibow kdrlcreert treated at | f1 14dpf_treated at 1.5dpf full trunk pcv cells 5mm | | treated at 1.5dpf | kdrl | na | x | x | na | \ | |
| 108 | 2021.2.21 flibow kdrlcreert treated at | f1 14dpf_treated at 1.5dpf full trunk pcv cells 5mm | | treated at 1.5dpf | kdrl | na | x | x | na | \ | |
| 109 | 2021.2.21 flibow kdrlcreert treated at | f1 14dpf_treated at 1.5dpf full trunk pcv cells 5mm | | treated at 1.5dpf | kdrl | na | x | x | na | \ | |
| 110 | 2021.2.21 flibow kdrlcreert treated at | f2 12dpf_treated at 1.5dpf close to the tail after da labeled | | treated at 1.5dpf | kdrl | na | na | x | x | \ | |
| 111 | 2021.2.21 flibow kdrlcreert treated at | f2 16dpf_treated at 1.5dpf all trunk 5.5mm | | treated at 1.5dpf | kdrl | na | na | x | x | \ | |
| 112 | 2021.2.21 flibow kdrlcreert treated at | f2 19dpf_treated at 1.5dpf anal pore 6mm | | treated at 1.5dpf | kdrl | na | na | x | x | \ | |
| 113 | 2021.2.21 flibow kdrlcreert treated at | f3 12dpf_treated at 1.5dpf close to the tail | | treated at 1.5dpf | kdrl | na | x | na | na | \ | |
| 114 | 2021.2.21 flibow kdrlcreert treated at | f3 16dpf_treated at 1.5dpf close to the tail | | treated at 1.5dpf | kdrl | na | x | na | na | \ | |
| 115 | 2021.2.21 flibow kdrlcreert treated at | f3 19dpf_treated at 1.5dpf close to the tail | | treated at 1.5dpf | kdrl | na | x | na | na | x | |
| 116 | 2021.2.21 flibow kdrlcreert treated at | f3 23dpf_treated at 1.5dpf close to the tail | | treated at 1.5dpf | kdrl | na | x | na | na | x | |
| 117 | | | | | | | | | | | |
| 118 | | | | | | | | | | | |
| 119 | 2021.2.21 flibow kdrlcreert treated at | f3 12dpf_treated at 1.5dpf anal pore to mid trunk | | treated at 1.5dpf | kdrl | na | na | x | na | \ | |
| 120 | 2021.2.21 flibow kdrlcreert treated at | f3 14dpf_treated at 1.5dpf all trunk_5.5mm | | treated at 1.5dpf | kdrl | na | na | x | na | \ | |
| 121 | 2021.2.21 flibow kdrlcreert treated at | f3 16dpf_treated at 1.5dpf all trunk_6mm | | treated at 1.5dpf | kdrl | na | na | x | na | \ | |
| 122 | 2021.2.21 flibow kdrlcreert treated at | f3 19dpf_treated at 1.5dpf mid trunk | | treated at 1.5dpf | kdrl | na | na | x | na | \ | |
| 123 | 2021.2.21 flibow kdrlcreert treated at | f3 23dpf_treated at 1.5 ù | | treated at 1.5dpf | kdrl | na | na | x | na | x | |
| 124 | 2021.2.21 flibow kdrlcreert treated at 1.5dpf tamoxifen 10um | | | treated at 1.5dpf | | | | | | | |
| 125 | | | | | | | | | | | |
| 126 | 2021.2.21 flibow kdrlcreert treated at | f4 12dpf_treated at 1.5dpf | | treated at 1.5dpf | kdrl | na | x | x | x | \ | |
| 127 | 2021.2.21 flibow kdrlcreert treated at | f4 14dpf_treated at 1.5dpf_5mm | | treated at 1.5dpf | kdrl | na | x | x | x | \ | |
| 128 | 2021.2.21 flibow kdrlcreert treated at | f4 16dpf_treated at 1.5dpf_5.2mm | | treated at 1.5dpf | kdrl | na | x | x | x | \ | |
| 129 | 2021.2.21 flibow kdrlcreert treated at | f4 19dpf_treated at 1.5dpf anal pore 7mm | | treated at 1.5dpf | kdrl | na | x | x | x | x | |
| | | f5 12dpf | | treated at 1.5dpf | kdrl | na | x | x | na | na | |
| | | f5 14dpf | | treated at 1.5dpf | kdrl | na | x | x | na | na | |
| | | f5 16dpf | | treated at 1.5dpf | kdrl | na | x | x | na | na | |
| | | f5 23dpf | | treated at 1.5dpf | kdrl | na | x | x | na | x | |
| 130 | 2020.11.19 flibow kdrlcreert treat at 3d | f1 flibow kdrlcreert treat at 3dpf for 24tamoxifen 14dpf zoo | | treated at 3dpf | kdrl | na | x | x | na | \ | |
| 131 | 2020.11.19 flibow kdrlcreert treat at 3d | f1 flibow kdrlcreert treat at 3dpf for 24tamoxifen 17dpf zoo | | treated at 3dpf | kdrl | na | x | x | na | x | |
| 132 | 2020.11.19 flibow kdrlcreert treat at 3dpf for 24h tamox | | | | | | | | | | |
| 133 | 2020.11.19 flibow kdrlcreert treat at 3d | f1 flibow kdrlcreert treat at 3dpf for 24tamoxifen 14dpf zoo | | treated at 3dpf | kdrl | na | x | x | x | \ | |
| 134 | 2020.11.19 flibow kdrlcreert treat at 3d | f1 flibow kdrlcreert treat at 3dpf for 24tamoxifen 17dpf zoo | | treated at 3dpf | kdrl | na | x | x | x | x | |
| 135 | 2020.11.19 flibow kdrlcreert treat at 3d | f5 flibow kdrlcreert treat at 3dpf for 24tamoxifen_14dpf all | | treated at 3dpf | kdrl | na | x | x | x | \ | |
| 136 | 2020.11.19 flibow kdrlcreert treat at 3d | f5 flibow kdrlcreert treat at 3dpf for 24tamoxif | | treated at 3dpf | kdrl | na | x | x | \ | x | |
| 137 | 2020.11.19 flibow kdrlcreert treat at 3d | f5 flibow kdrlcreert treat at 3dpf for 24tamoxifen 2 | | treated at 3dpf | kdrl | na | | | \ | x | |
| | | | | | | na | | | | | |
| 138 | 2020.11.19 flibow kdrlcreert treat at 3d | f5 flibow kdrlcreert treat at 3dpf for 24tamoxifen_14dpf zo | | treated at 3dpf | kdrl | na | x | x | na | \ | |
| 139 | 2020.11.19 flibow kdrlcreert treat at 3d | f5 flibow kdrlcreert treat at 3dpf for 24tamoxifen_14dpf zo | | treated at 3dpf | kdrl | na | x | x | na | x | |
| 140 | 2020.11.19 flibow kdrlcreert treat at 3dpf for 24h tamox | | | treated at 3dpf | kdrl | na | x | x | na | x | |
| 141 | 2020.11.30 flibow hscreert treat 3dpf 1 | f1_flibow_hscreert treated at 3dpf 10um tail 14dpf | | treated at 3dpf | hs | x | x | x | x | \ | |
| 142 | 2020.11.30 flibow hscreert treat 3dpf 1 | f1_flibow_hscreert treated at 3dpf 10um blu inters 20dpf | | treated at 3dpf | hs | x | x | x | x | x | |
| 143 | 2020.11.30 flibow hscreert treat 3dpf 1 | f2_flibow_hscreert treated at 3dpf 10um blu da 14dpf 20 | | treated at 3dpf | hs | x | x | x | x | \ | |
| 144 | 2020.11.30 flibow hscreert treat 3dpf 1 | f3_flibow_hscreert treated at 3dpf 10um vc 20dpf_2023-1 | | treated at 3dpf | hs | na | x | na | na | x | |
| | | f3_flibow_hscreert treated at 3dpf 10um vc 23dpf_2023-1 | | treated at 3dpf | hs | na | x | na | na | x | |
| | 2020.29.10 flibow hscreert treated at 24hpf 10 min hs 30 min tamox.czi | | | | | | | | | | |
| 145 | 2020.29.10 flibow hscreert treated at 2 | f1_flibow hscreert treated at 24hpf 10 min hs 30 min tamox | | treated at 1.5dpf | hs | na | x | x | x | \ | circulating cells labeled |
| 146 | 2020.29.10 flibow hscreert treated at 2 | f2_flibow hscreert treated at 24hpf 10 min hs 30 min tamox | | treated at 1.5dpf | hs | na | x | na | na | \ | circulating cells labeled |
| 157 | 2020.29.10 flibow hscreert treated at 2 | f3_flibow hscreert treated at 24hpf 10 min hs 30 min tamox | | treated at 1.5dpf | hs | na | x | x | na | \ | circulating cells labeled |
| 148 | 2020.29.10 flibow hscreert treated at 2 | f4_flibow hscreert treated at 24hpf 10 min hs 30 min tamox | | treated at 1.5dpf | hs | na | x | x | na | \ | circulating cells labeled |
| | 2021.3.29 flibow kdrlcre treated at 3dpf 10um | | | | | | | | | | |
| 149 | 2021.3.29 flibow kdrlcre treated at 3dp | f1 treated at 3dpf 10 um 12dpf full trunk 5mm | | treated at 3dpf | kdrl | na | x | x | na | \ | |
| 150 | 2021.3.29 flibow kdrlcre treated at 3dp | f2 treated at 3dpf 10 um 12dpf full trunk 5mm | | treated at 3dpf | kdrl | na | x | x | x | \ | |
| 152 | 2021.3.29 flibow kdrlcre treated at 3dp | f3 treated at 3dpf 10 um 12dpf full trunk 5mm | | treated at 3dpf | kdrl | na | x | x | x | \ | |
| 153 | 2021.3.29 flibow kdrlcre treated at 3dp | f4_treated at 3dpf 10 um 12dpf full trunk 5mm | | treated at 3dpf | kdrl | na | x | x | x | \ | |
| 154 | 2021.3.29 flibow kdrlcre treated at 3dp | f5_treated at 3dpf 10 um 12dpf full trunk 4.5mm | | treated at 3dpf | kdrl | na | x | x | x | \ | |
| 155 | 2021.3.29 flibow kdrlcre treated at 3dp | f6_treated at 3dpf 10 um 12dpf full trunk 4.5mm | | treated at 3dpf | kdrl | na | x | x | x | \ | |
| 156 | 2021.3.29 flibow kdrlcre treated at 3dp | f7_treated at 3dpf 10 um 12dpf full trunk 4.5mm | | treated at 3dpf | kdrl | na | x | x | x | \ | |
| | | | | | kdrl | | | | | | |
| 157 | 2021.1.17 flibow kdrlcreert treated at 2 | f1_13dpf all trunk_a_f1_13dpf all trunk image | | treated at 2dpf | kdrl | na | x | x | na | \ | |
| 158 | 2021.1.17 flibow kdrlcreert treated at 2 | f1_16dpf all trunk_a_f1_16dpf all trunk image | | treated at 2dpf | kdrl | na | x | x | na | \ | |
| | | | | | | na | | | | | |
| 159 | 2021.1.17 flibow kdrlcreert treated at 2 | f2_13dpf anal pore1 cell | | treated at 2dpf | kdrl | na | na | x | na | \ | |
| 160 | 2021.1.17 flibow kdrlcreert treated at 2 | f2_16dpf anal pore1 cell | | treated at 2dpf | kdrl | na | na | x | na | \ | |
| 161 | 2021.1.17 flibow kdrlcreert treated at 2dpf x 24h 10um | | | | | | | | | | |
| | | f3_13dpf full trunk | | treated at 2dpf | kdrl | na | x | x | x | \ | |
| 162 | 2021.1.17 flibow kdrlcreert treated at 2 | f3_16dpf full trunk not good | | treated at 2dpf | kdrl | na | x | x | x | \ | |
| 163 | 2021.1.17 flibow kdrlcreert treated at 2 | f3 all trunk 20dpf | | treated at 2dpf | kdrl | na | x | x | x | x | |
| 164 | 2021.1.17 flibow kdrlcreert treated at 2 | f4_13dpf anal pore 1 cell label | | treated at 2dpf | kdrl | na | x | x | na | \ | |
| 165 | 2021.1.17 flibow kdrlcreert treated at 2 | f4_16dpf anal pore 1 cell label | | treated at 2dpf | kdrl | na | x | x | na | x | |
| 166 | 2021.1.17 flibow kdrlcreert treated at 2 | f4 mid trunk inters 20dpf | | treated at 2dpf | kdrl | na | x | x | na | x | |
| | | | | | | na | | | | | |
| 167 | 2021.1.17 flibow kdrlcreert treated at 2 | f5_13dpf blu inters tail | | treated at 2dpf | kdrl | na | x | x | na | \ | |
| 168 | 2021.1.17 flibow kdrlcreert treated at 2 | f6_13dpfall trunk l_a_f6_13dpfall trunk l_image 1_a_f6_ | | treated at 2dpf | kdrl | na | x | x | x | \ | |
| 169 | 2021.1.17 flibow kdrlcreert treated at 2 | f7 14 dpf all trunk | | treated at 2dpf | kdrl | na | x | x | x | \ | |
| 170 | 2021.1.17 flibow kdrlcreert treated at 2 | f7 16 dpf all trunk | | treated at 2dpf | kdrl | na | x | x | x | \ | |
| 171 | 2021.1.17 flibow kdrlcreert treated at 2 | f7 all trunk_20dpf | | treated at 2dpf | kdrl | na | x | x | x | x | |
| 172 | 2021.1.17 flibow kdrlcreert treated at 2 | f8 only blu loop mid trunk good 4.5mm 14dpf | | treated at 2dpf | kdrl | na | na | x | na | \ | |
| 173 | 2021.1.17 flibow kdrlcreert treated at 2 | f8 only blu loop mid trunk good 6mm 16dpf | | treated at 2dpf | kdrl | na | na | x | na | \ | |
| 174 | 2021.1.17 flibow kdrlcreert treated at 2 | f8 only blu loop mid trunk good 8.5mm 20dpf | | treated at 2dpf | kdrl | na | na | x | na | x | |
| 175 | 2021.1.17 flibow kdrlcreert treated at 2 | f8 only blu loop mid trunk good 10mm 26dpf | | treated at 2dpf | kdrl | na | na | x | na | x | |
| 176 | 2021.1.17 flibow kdrlcreert treated at 2 | f9 14 dpf | | treatedat 2dpf | kdrl | na | x | x | x | \ | |
| 177 | 2021.1.17 flibow kdrlcreert treated at 2 | f9 16 dpf full trunk | | treatedat 2dpf | kdrl | na | x | x | x | \ | |
| 178 | 2021.1.17 flibow kdrlcreert treated at 2 | f10 14 dpf anal pore | | treated at 2dpf | kdrl | na | x | x | x | \ | |
| 179 | 2021.1.17 flibow kdrlcreert treated at 2 | f10 16 dpf full trunk | | treated at 2dpf | kdrl | na | x | x | x | \ | |
| 180 | 2021.1.17 flibow kdrlcreert treated at 2 | f10 full trunk_20dpf | | treated at 2dpf | kdrl | na | x | x | x | x | |
| 181 | 2021.1.17 flibow kdrlcreert treated at 2 | f11 14 dpf all trunk_a_f11 14 dpf all trunk_image | | treated at 2dpf | kdrl | na | x | x | x | \ | |
| 182 | 2021.1.17 flibow kdrlcreert treated at 2 | f11 16 dpf all trunk xrudra | | treated at 2dpf | kdrl | na | x | x | x | \ | |
| 183 | 2022.10.3 flibow kdrlcreert_treated 3d | f4 flibow kdrlcreert treated 3dpf 24h 14dpf | | treated at 3dpf | kdrl | na | x | x | x | \ | |
| 184 | 2022.10.3 flibow kdrlcreert_treated 3d | f4 flibow kdrlcreert treated 3dpf 24h 17dpf | | treated at 3dpf | kdrl | na | x | x | x | \ | |
| 185 | 2022.10.3 flibow kdrlcreert_treated 3d | f5 flibow kdrlcreert treated 3dpf 24h 14dpf | | treated at 3dpf | Kdrl | na | x | x | na | \ | |
| 186 | 2022.10.3 flibow kdrlcreert_treated 3d | f5 flibow kdrlcreert treated 3dpf 24h 23dpf | | treated at 3dpf | Kdrl | na | x | x | na | \ | |
| 187 | 2022.10.3 flibow kdrlcreert_treated 3d | f5 flibow kdrlcreert treated 3dpf 24h 34dp | | treated at 3dpf | Kdrl | na | x | x | na | x | |

51
